# Ecological predictors of organelle genome evolution: Phylogenetic correlations with taxonomically broad, sparse, unsystematized data

**DOI:** 10.1101/2023.08.11.553003

**Authors:** Konstantinos Giannakis, Luke Richards, Iain G. Johnston

## Abstract

Comparative analysis of variables across phylogenetically linked observations can reveal mechanisms and insights in evolutionary biology. As the taxonomic breadth of the sample of interest increases, challenges of data sparsity, poor phylogenetic resolution, and complicated evolutionary dynamics emerge. Here, we investigate a cross-eukaryotic question where all these problems exist: which organismal ecology features are correlated with gene retention in mitochondrial and chloroplast DNA (organelle DNA or oDNA). Through a wide palette of synthetic control studies, we characterize the specificity and sensitivity of a collection of parametric and non-parametric phylogenetic comparative approaches to identify relationships in the face of such sparse and awkward datasets. We combine and curate ecological data coupled to oDNA genome information across eukaryotes, including a new semi-automated approach for gathering data on organismal traits from less systematized open-access resources including encyclopedia articles on species and taxa. Combining this unique dataset with our benchmarked comparative approaches, we confirm support for several known links between organismal ecology and organelle gene retention, identify several previously unidentified relationships constituting possible ecological contributors to oDNA genome evolution, and provide support for a recently hypothesized link between environmental demand and oDNA retention. We, with caution, discuss the implications of these findings for organelle evolution and of this pipeline for broad comparative analyses in other fields.

## Introduction

The statistical investigation of variables that are correlated across biological species can help reveal evolutionary and organismal mechanisms and relationships (Paradis 2014). In such investigation, controlling for the relatedness of species is essential to avoid false positive and negative results. For example, there are thousands of eukaryotic species which have both fur and four limbs. But these do not constitute thousands of independent samples supporting a correlation between these variables, because they inherited those properties from a common mammalian ancestor. A wealth of approaches exists for comparative analyses accounting for such phylogenetic structure. Early milestones in addressing this questions were Felsenstein’s independent contrasts approach (Felsenstein 1985) and Grafen’s phylogenetic regression with generalized least squares (Grafen 1989), originally developed with an explicit or implicit picture of continuously varying characters. More recently, the range of approaches has dramatically expanded to offer statistically powerful methods for specific circumstances (Paradis 2014). For questions of variables that do not follow the continuously-varying picture, these include a development by Pagel for binary characters (Pagel 1994), and phylogenetically-embedded linear and generalized linear models for different response variable structures (Ives and Garland 2010; Paradis and Claude 2002). (Maddison and FitzJohn 2015) discuss how several well-principled approaches still suffer in practice from the problem of pseudoreplication, where identical values of a discrete trait can arise within a clade by common descent rather than through a “generative” correlation. (Maddison 2000) proposes a different class of approach where phylogenetically independent sets of observations are identified and analysed together; (Maddison and FitzJohn 2015) discuss an approach where such independent subsets of the phylogeny are identified before being subjected to, for example, Pagel’s approach (Pagel 1994). Careful treatments and commentaries emphasise that the statistical power and effect of correcting for phylogenetic correlations depend on the specifics of the problem (Revell 2010; Rohle 2006), and that detailed investigation of a method’s performance on synthetic data can make interpretation clearer. A common theme behind these discussions is the necessity of comparing summary statistics and hypothesis testing outcomes to simulations of null and alternative hypotheses, to characterize the influence of any assumption breaking or other challenges to method applicability.

Here, we investigate a particular evolutionary question where many common comparative assumptions are broken: what species-specific features are linked to the retention of genes in organelle DNA (oDNA)? Across eukaryotes, gene content in mitochondrial DNA (mtDNA) and plastid DNA (ptDNA) has been dramatically reduced since the endosymbiotic events that created the organelles, and now varies dramatically across species (D. R. Smith and Keeling 2015; Giannakis et al. 2022; Johnston and Williams 2016; Mohanta et al. 2020; Roger, Muñoz-Gómez, and Kamikawa 2017; Keeling 2010; Johnston and Burgstaller 2019; Janouškovec et al. 2017). The genes retained in oDNA are of central importance in bioenergetics and metabolism, with impact from human diseases (Stewart and Chinnery 2015; Taylor and Turnbull 2005; Wallace and Chalkia 2013) to crop production (Chen and Liu 2014; Mackenzie 2010; Havey 2004). The question of why particular *genes* are retained across species has been studied for some time: hypotheses include the favouring of genes that are hard to import to the organelle (von Heijne 1986; Björkholm et al. 2015), allow direct local control of organelle function (Allen 2015; Allen and Martin 2016), are energetically favourable to retain (Kelly 2021), and many more, reviewed and quantitatively compared in (Giannakis et al. 2022; Johnston and Williams 2016). But the dual question of why particular *species* retain a given number of genes has received less attention. Some specific hypotheses have been proposed: parasitic organisms are more free to lose or transfer control of their bioenergetic organelles (Hjort et al. 2010; Keeling 2010; D. R. Smith and Keeling 2015), gene transfer is more beneficial in plants with self-fertilisation and clonal modes of reproduction (Brandvain, Barker, and Wade 2007; Brandvain and Wade 2009), and different energetic economics across taxa may help shape oDNA gene profiles (Kelly 2021). However, with these exceptions, the general reasons why different species retain different organelle gene counts remain open.

Recent modelling work has suggested that a species’ environment may be responsible for some of this diversity (García-Pascual, Nordbotten, and Johnston 2022). Specifically, the retention of more oDNA genes is predicted to be beneficial in organisms whose environments provide strong, oscillating metabolic or bioenergetic demands (for example, diurnal or tidal oscillations). In such environments, oDNA gene retention ensures that individual organelles can respond to changing demands more rapidly and individually than if essential genes are transferred to the nucleus – following the colocation for redox regulation (CoRR) hypothesis (Allen 2015; Allen and Martin 2016). Organisms that exist in more stable, less demanding environments (or can move away from challenges) are predicted to have less need for rapid, local organelle responses, and thus retain fewer oDNA genes. This theory captures some broad observations: intracellular parasites typically retain very few oDNA genes (Hjort et al. 2010; Keeling 2010); fungi and motile metazoans (less exposed to, or capable of moving from, fluctuating environmental conditions) retain more; plants (subject to diurnal fluctuations) more still (Johnston 2019); and algae and other species in intertidal and estuarine conditions often retain more yet (Keeling 2010). We wanted to ask whether more specific information on species ecology and environments could help quantitatively support this hypothesis.

However, the nature of this question poses several challenges to a data-driven comparative analysis. In a sense, the simplest case for phylogenetic comparative work is two continuously-varying characters, one predictor and one response, that are perfectly observed and evolve according to Brownian motion on a perfectly known (and balanced) phylogeny (Symonds and Blomberg 2014; Felsenstein 1985). Our system departs from this ideal in several ways:

- The predictor is an ecological factor (often binary) and the response is an ordinal (gene count);
- Evolution proceeds by monotonic decrease of the response variable, at a rate that may depend on the predictor;
- The branch lengths of the phylogeny are not known in general;
- Some clades are both highly overrepresented in the dataset and have very similar response values (for example, metazoans, which form the majority of mtDNA samples and all have almost identical mtDNA gene profiles);
- The observations of the predictor on the phylogeny’s tips are sparse: a positive observation typically corresponds to a positive value, but a negative observation may correspond either to a negative value or to an absent observation.

We therefore proceed by establishing a set of synthetic control cases where the performance of comparative methods is tested on simulated systems reflecting these complications (Revell 2010). The insight from these controls, coupled with curation of ecological and oDNA sequence data, allows us to establish and interpret the test cases: comparative analysis of oDNA gene counts and variables describing organismal ecology and environments across eukaryotes.

## Methods

### Simulating organelle-like evolution and data sparsity

We built a series of synthetic models where predictor and response co-evolved on artificially constructed phylogenetic trees (Fig. 1). These trees either had uniform branch lengths or branch lengths determined by a simulated birth-death process with different parameterisations (Stadler 2011; Paradis and Schliep 2019). The evolutionary rule for constructing descendant state (*X_d_*, *Y_d_*) from ancestor state (*X_a_*, *Y_a_*), given a branch of length *t_a,d_*linking the two, was

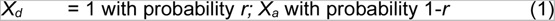

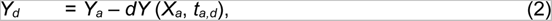

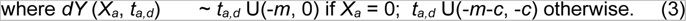

**Figure 1.**
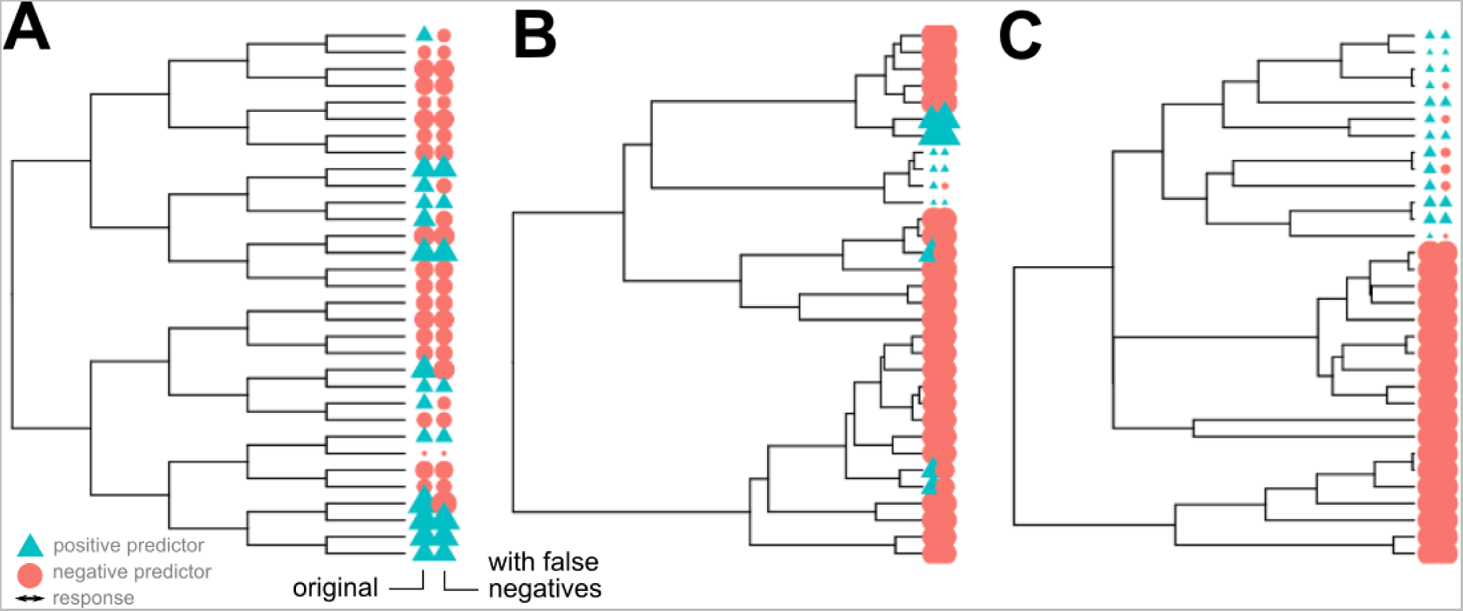
Example simulated phylogenies in the control studies. In each case the tip labels give the true states (left) and observed states after false negative observations (right). Colour gives predictor value (red circles negative, blue triangles positive); size gives response value. (A) A symmetric, balanced tree with no influence of predictor on response. (B, C) Birth-death trees with different death parameters, with strong influence of predictor on response.

In other words, each ancestor-descendant step has a probability *r* of changing the descendant to a positive *X* value. The descendant’s *Y* value is inherited from its ancestor, with a random reduction *dY* of characteristic size *m* scaled by the branch length, and shifted to higher magnitudes if the ancestor’s X value is nonzero. Any negative *Y* values that emerged under Eqns. 1-3 are set to zero. To choose a particular metaphor, parasitism (*X*=1) evolves at random through the phylogeny, and increases the rate of mtDNA gene loss when it appears.

After simulating evolution, each tip on the phylogeny had a pair of values (*X*, *Y*). Those tips with *X* = 1 were assigned an “observed” value *X* = 0 with probability *p*. This false negative observation modelling is designed to mimic the false negatives which are known to be prevalent in several of our datasets, where records of organismal traits are sparse: a positive observation likely corresponds to a true positive, but a negative observation could correspond to a true negative or incomplete observations.

The parameters we varied were *r*, the probability of the predictor changing value along a given branch; *c*, the influence of the *X*=1 case on the rate of *Y* evolution; and *p*, the probability with which a true *X*=1 value led to an *X*=0 observation. We also varied the size of the tree *n* and *ν*, the death rate parameter in the birth-death process that generated random phylogenies.

### Phylogenetic generalized least squares and (generalized) linear models

We used R packages *ape* (Paradis and Schliep 2019) and *nlme* (DebRoy 2006) to implement phylogenetic generalized least squares (PGLS) (Revell and Harmon 2022; Grafen 1989; Symonds and Blomberg 2014). Specifically, we used a range of evolutionary models (Brownian motion (Felsenstein 1985; Martins and Hansen 1997), Grafen’s method (Grafen 1989), scaling covariances by Pagel’s *λ* (Freckleton, Harvey, and Pagel 2002; Pagel 1999), and the Martins-Hansen approach of exponentially decaying covariance over evolutionary time (Martins and Hansen 1997)) to estimate the covariance matrix linking tip observations on a phylogeny, and used this correlation structure in a generalized least squares fit relating our predictor and response variables. P-values were provided based on a chi-squared null distribution (Susko 2003), which in turn assumes normality of residuals, which is generally not the case for our data. Hence, we test the conclusions from this analysis against the simulated systems above to assess the validity of this interpretation. For phylogenetic generalized linear models (PGLM) we used the *phylolm* package (Tung Ho and Ané 2014) for Poisson regression accounting for phylogenetic correlation with generalized estimating equations (Paradis and Claude 2002); p-values here again are dependent on some parametric assumptions, so we refer to tests against simulation for interpretation as in the text. We also used *phylolm* in parallel to perform phylogenetic linear modelling (PLM).

### Within-family comparisons

Our non-parameteric approach for comparing relatives follows the broad idea in (Maddison 2000). We first assign *X* and *Y* values to all internal nodes in the phylogeny. Predictor *X* is set to positive for an ancestor if all its descendants have positive values, otherwise the ancestor is set to negative. Response values *Y* for an ancestor are set to the mean of its descendants. We then identify sets of siblings (among internal nodes as well as tips) which differ in their *X* values. For each, we record *Y^+^* and *Y^−^*, respectively the mean response value across the positive- and negative-predictor-valued siblings. The list of (*Y^+^* − *Y^−^*) values is recorded across the sets of siblings. Either a Wilcoxon rank-sum test or bootstrap resampling is then used to test the median of this list against a null hypothesis of zero (siblings with positive and negative predictor values have no difference in response values).

### Accounting for clade influence

Much variability in oDNA gene count occurs at the level of supergroups and kingdoms (Janouškovec et al. 2017; Giannakis et al. 2022). In a parallel approach attempting to account for this, we also used several approaches to “block” different eukaryotic clades as sources of oDNA gene count variability and analyse the remaining variance. We used linear mixed models and Poisson generalized linear mixed models (LMMs and GLMMs) with random effects on either intercepts alone, or intercepts and slopes (selected via Akaike Information Criterion, AIC), associated with eukaryotic clade. We also took a non-parametric approach using the Scheirer-Ray-Hare (SRH) test, which resembles a generalization of the better-known Kruskal-Wallis test to allow for two factors and replicated observations (Scheirer, Ray, and Hare 1976). Specifically, the first assigned branch after the common eukaryotic ancestor from NCBI’s Common Tree was used as the blocking variable, and the feature of interest was used as the group variable. If a particular feature was only found to take different values in a single clade, the Kruskal-Wallis test was used within that single clade to provide an analogous result. The SRH test was implemented using the *rcompanion* package (Mangiafico 2020). Finally, we attempted to follow the philosophy of (Maddison and FitzJohn 2015), where we independently normalized the gene counts within a clade by subtracting that clade’s mean count from every observation, before analysing with PLM as above.

### Curation of organismal properties from databases

The trait dataset used in our analysis was assembled using a combination of automated and manual methods. Species names and genetic information (such as organellar genome size, GC content, and taxonomical information) were collected via NCBI’s Organelle database (O’Leary et al. 2016) and processed according to the pipeline in (Giannakis et al. 2022). After curation, we retained 9296 mtDNA sequences (8835 of which are metazoans) and 4264 ptDNA sequences. The majority of the ecological trait entries were acquired via Encyclopedia of Life (Parr et al. 2014) through individual queries for traits of interest, and then cross-referenced with some domain-specific databases (like GloBI (Poelen, Simons, and Mungall 2014), but also others, see following references). Encyclopedia of Life is an open encyclopedia that provides a catalogue with plenty of traits and taxonomic information on organisms, which consists of contributions and curation by experts and organisations. Palm data were collected through (Kissling et al. 2019). Organismal habitats and traits related to cross-species interactions (including parasitism) were taken from (Cohen et al. 2020), a large ecological scale survey on species interactions. For the majority of plant data, ecological traits and characters were taken from Encyclopedia of Life and crosschecked with the TRY database (2018 update) (Kattge et al. 2020).

For the cross-eukaryote data, NCBI’s Common Taxonomy Tool was used to estimate phylogenetic topology (Federhen 2012). More specific phylogenetic information was obtained through the U.PhyloMaker R package (Jin and Qian 2023; 2022) which uses the Plant megaphylogeny found in (Jin and Qian 2022) comprised by phylogenies from (S. A. Smith and Brown 2018) and (Zanne et al. 2014). The plant phylogeny assembled in (Jin and Qian 2022) and sourced from (S. A. Smith and Brown 2018; Zanne et al. 2014) gave us a sufficient number of matches (up to the level of genus) to our list of species (120 for the species with mitochondrial gene count and 3851 for plastid) and we were able to test different scenaria.

Besides the automated retrieval of trait values from the above resources via custom R and Python scripts, manual insertions and corrections took place over individual observations, either through parsing Wikipedia pages (see below) or via ongoing literature reading and personal communications. Data were processed and assembled into two major datasets (one for each organelle) using custom R scripts.

### Semi-automated curation of organismal properties from less systematized sources

The approach assumes that an XML dump of articles (for example, Wikipedia pages, PubMed abstracts, or NCBI entries) corresponding to a set of taxa of interest can be obtained (for example, via Wikipedia’s Special:Export API (https://en.wikipedia.org/wiki/Special:Export). A custom Python script then searches for a given regular expression pattern in such an XML file, and produces an HTML file designed to assist manual parsing of all the entries matching that pattern. The HTML page consists of the blocks of text surrounding each pattern match, labelled by page name. Hyperlinks and other interesting text are accompanied by checkboxes, which when clicked populate a text box on the right-hand side of the page with the text immediately before the checkbox. At the bottom of the HTML page there is a summary button which compiles all the text from other boxes into a single comma-separated list.

The user can quickly check any examples where a given element of text (for example, a species or taxon name) genuinely corresponds to the feature of interest, then compile all these positive cases into a summary list. Regular expressions can be arbitrarily complex (for example, to identify single-celled organisms we found */[Uu]nicell|[Uu]ni-cell|[Ss]ingle-cell/* to give useful results).

### Software references

We used R (R Core Team and Team 2022) with packages *ape* (Paradis and Schliep 2019), *phytools* (Revell 2012), *phylolm* (Tung Ho and Ané 2014), and *phangorn* (Schliep 2011) for phylogenetic analysis, *lme4* (Bates et al. 2015), *nlme* (DebRoy 2006) and *rcompanion* (Mangiafico 2020) for statistical analysis, and *ggplot2* (Wickham 2011), *beeswarm* (Eklund 2016), *ggrepel* (Slowikowski 2021), *gridExtra* (Auguie and Antonov 2017), *ggtree* (Yu et al. 2017), and *ggtreeExtra* (Xu et al. 2021) for plotting. ODNA profiles were taken from the pipeline in (Giannakis et al. 2022). All software and data for the phylogenetic analysis is available at https://github.com/StochasticBiology/comparative-odna.

## Results

### Sensitivity and specificity of comparative phylogenetic methods with organelle-like evolution and varying data properties

We sought to understand the impact of all the above issues on the performance of comparative approach seeking correlations between predictor and response. We simulated a range of synthetic evolutionary systems on different tree topologies, involving a response variable decreasing at a rate that may depend on a predictor variable that irreversibly changes state at random. Observations of the predictor variable were occluded with a false negative probability (see Methods; Fig. 1). This setup is designed to match the complications in organelle genome analysis.

Because of its flexibility, we began with phylogenetic generalized least squares (PGLS) using a Brownian evolution model to estimate species correlations (Symonds and Blomberg 2014; Grafen 1989). We also explored other approaches for investigating correlations while accounting, or not, for phylogenetic coupling (Methods; Supplementary Fig. 1). We explored the use of different evolutionary models to estimate the covariance matrix linking observations (as well as the naïve case with no accounting for phylogenetic relationships). In parallel with PGLS, we tested an implementation of phylogenetic linear modelling (PLM) also using the Brownian evolution model (Tung Ho and Ané 2014), and phylogenetic generalized linear modelling (PGLM) with a Poisson response distribution via generalized estimating equations (Tung Ho and Ané 2014; Paradis and Claude 2002). We also used a nonparametric pipeline based on comparisons of related species or clades with different values of the predictor without invoking a particular evolutionary model, following the principle of (Maddison 2000). Finally, we explored the effects of considering or neglecting branch length information with different evolutionary models for covariance structure. We assessed the specificity and sensitivity of these methods as tree size, tree topology, false negative observation probability, magnitude of predictor-response relationship, and prevalence of positive predictor values were varied.

The naive approach neglecting phylogenetic structure and treating each species as an independent sample led to, as expected, a substantial false positive rate (as correlation by shared descent was not considered). The nonparametric approach performed poorly, with low statistical power, although false positives were limited. PGLS and PGLM, on the other hand, performed well at detecting extant links between variables, even with substantial departures from the idealized case above (Supplementary Figs. 1–3). An example is shown in Fig. 2. Not unexpectedly, the PLM approach behaved almost identically to the PGLS approach (Fig. 2, Supplementary Fig. 1).

**Figure 2.**
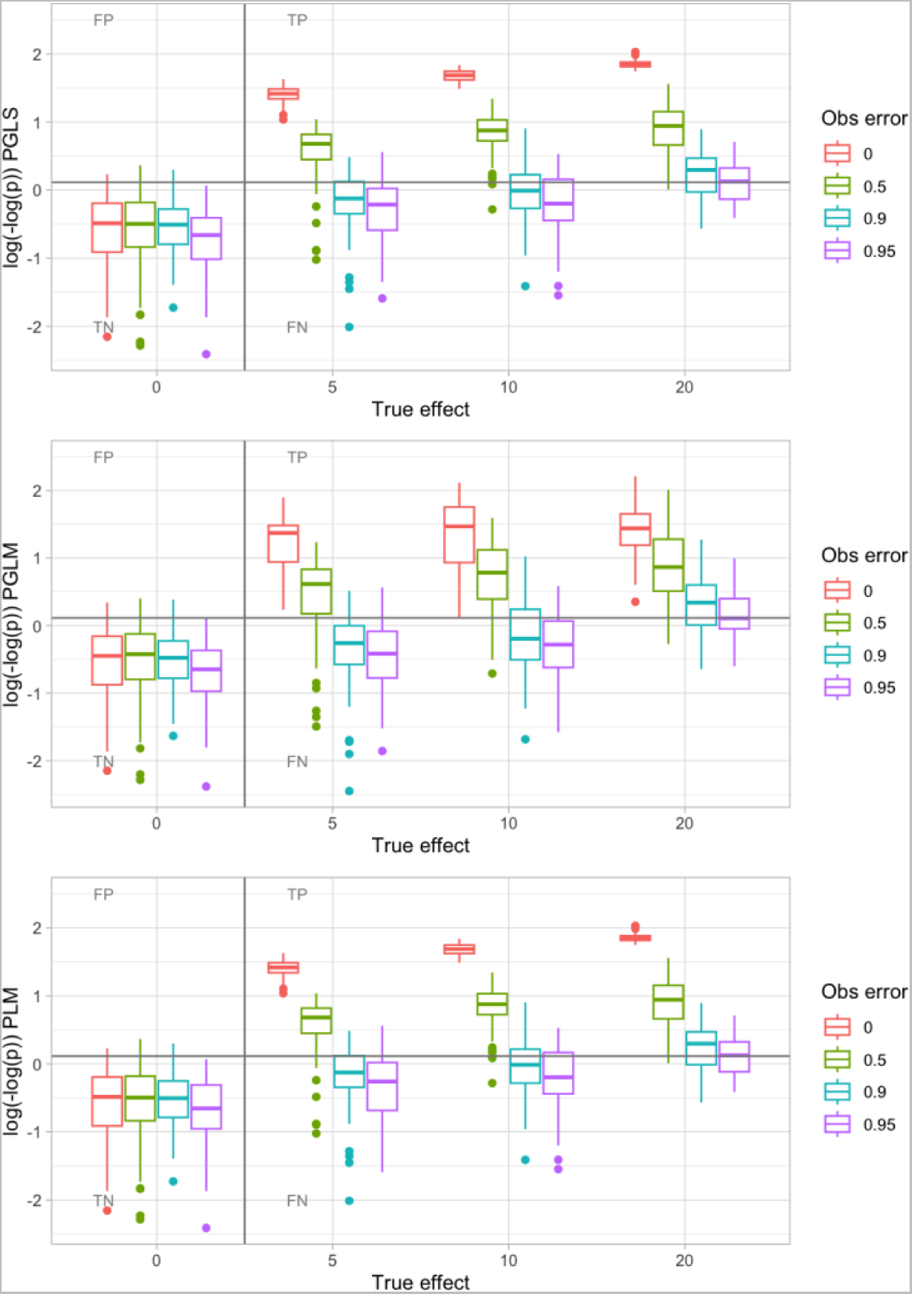
Example of PGLS, PLM, and PGLM sensitivity-specificity investigation. Evolutionary dynamics were simulated on a tree with 256 leaves, with differing true effect *c* linking predictor value and response evolution. Observations of the predictor value were occluded with an observation error parameter giving the probability that a positive value is observed as a negative. PGLS with Brownian correlation structure rarely gives false positive (FP) correlations and has substantial power to detect true positive (TP) correlations even for high observation error probabilities; PLM is almost identical in performance. PGLM likewise limits FP and retains power to detect TP, although the spread of p-values reported by PGLM for positive cases is rather broader. Here, an average of 8 evolutionary events innovated a positive predictor value; the effects of other simulation parameters and methods are shown in Supp Figs. 1-3.

Our interpretation of this perhaps surprisingly good performance for these parametric approaches is that although the evolutionary dynamics in our system are rather different from the ideal case (for example, continuous traits evolving according to Brownian motion), the latter still suffices to provide a reasonable estimate of the expected correlations between different species. Although the residual structure from our GLS model fit is not (cannot be) perfectly normal, the approach is general enough to allow informative followup analysis. The one-way occlusion of observations (positives may be observed as negative, but not the other way round) means that signal is preserved more than a completely random observation setup would allow.

One disadvantage of the PGLS approach is its relative computational intensity. For phylogenies of the size involved with our organelle data (over 9000 mtDNA sequences and 4000 ptDNA sequences), the construction and use of an *N* x *N* covariance matrix (Martins and Hansen 1997) becomes quite a computational challenge (taking several core days to analyse dozens of predictors). The PLM and PGLM approaches of estimating correlations using tree structure and generalized estimating equations (Paradis and Claude 2002; Tung Ho and Ané 2014) are much faster (taking only core seconds for the same analysis), as is the relative-comparison approach of recursively labelling a tree then seeking diverse siblings.

### Semi-automated pipeline for labelling organismal traits from unsystematized data sources

Encouraged by the potential of this comparative approach to detect correlations in the face of awkward and imperfectly observed data, we next attempted to gather information on organismal traits that could be linked to oDNA gene retention. Many of our traits of interest are not systematically assigned across a full list of eukaryotic species in a dedicated database (see Methods). However, less formalized online sources contain a substantial amount of information about some traits. Wikipedia (https://www.wikipedia.org/), for example, contains articles about many taxonomic groups and individual species, which often describe habitats, free-living vs parasitic lifestyles, multicellular vs unicellular physiology, and so on. However, fully automated methods to extract this information were challenging to construct. The huge variety of phrasings of this information and difficulties in automatically resolving ambiguous text suggested that a manual curation would be necessary. But reading (for example) the Wikipedia page for thousands of entries would be a tremendous investment of time.

We briefly explored using ChatGPT (https://chat.openai.com/chat) to gather organismal traits. This was unsuccessful. ChatGPT readily fabricated information about traits in both taxonomic groups and individual species. It was unable to either reliably assign traits to entities or to produce list of entities with a given trait.

To make progress, we constructed a pipeline to semi-automate the process of gathering information on a given trait from a source like Wikipedia (Methods). Illustrated in Fig. 3, this pipeline allows a user to provide a list of species and taxa to query, then returns every instance in the data source where an entry associated with elements of that list contain a term of interest. This set can either be immediately used or, as shown in Fig. 3, manually parsed to ensure reported entries genuinely correspond to the term of interest. We used this pipeline to gather information on several traits for our oDNA datasets: parasitism, extremophilia, multicellularity, sessility, flagellated physiology, and different photosynthetic modes, but we anticipate that its use could be much more general.

**Figure 3.**
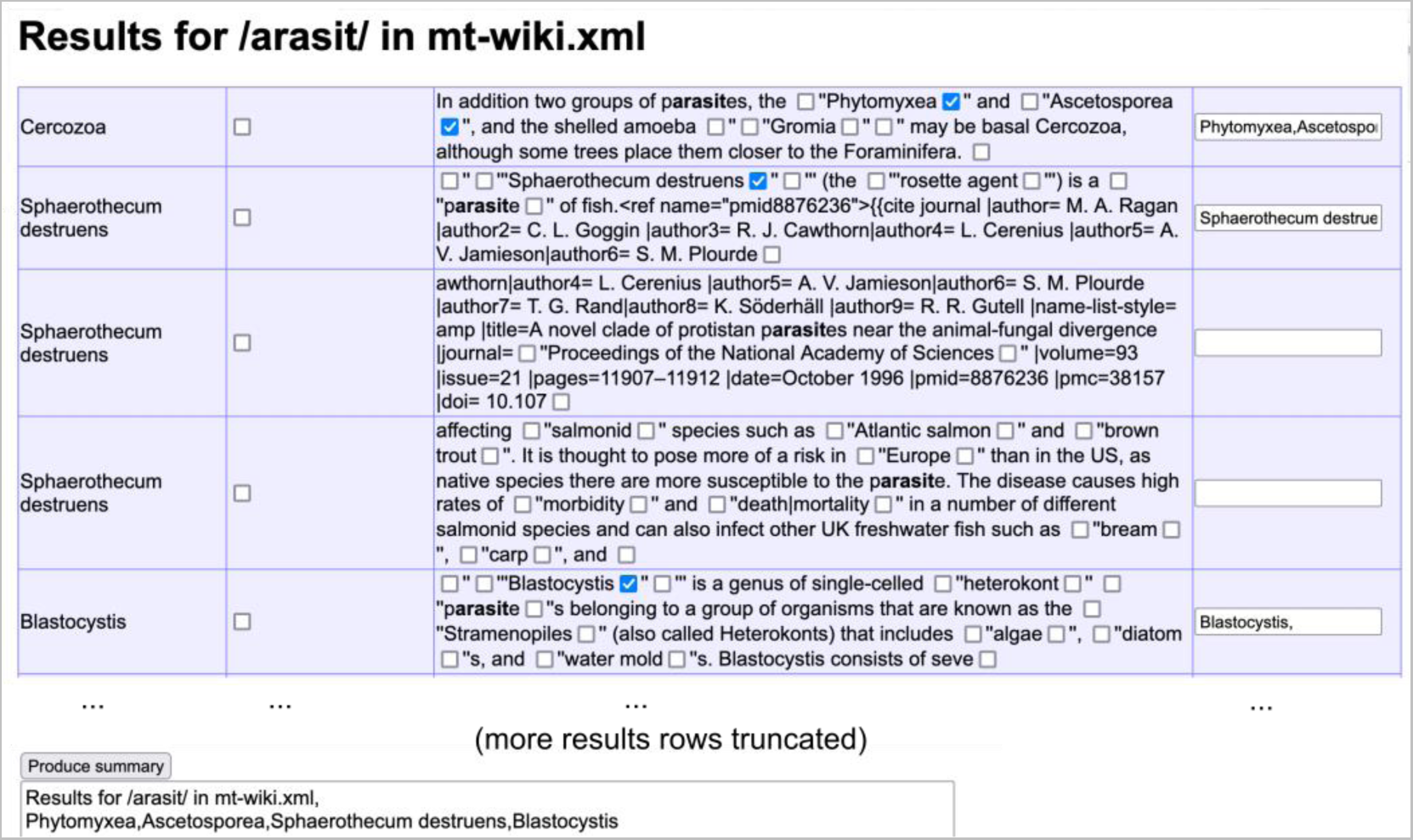
Screenshot of user interface for semi-automated extraction of organismal traits from online encyclopedia content. A custom script seeks a regular expression associated with a trait (here */arasit/*, designed to match [Pp]arasit[ic/ism/etc]) in the corpus of Wikipedia articles describing species and taxa within our phylogeny of interest. The text surrounding each instance of the expression is reported, with check boxes allowing the selection of hyperlinked terms that are manually deemed to match the trait (some examples demonstrated) – which are then stored in boxes on the right, where terms can also be manually entered. An example set of entries is shown in the figure; this query returned several hundred more. After parsing entries (many truncated here), a summary button creates a comma-separated list of all positively-identified or entered terms.

### Features correlated with mitochondrial and chloroplast gene counts across eukaryotes

Combining the above control studies, comparative workflow, and data-gathering pipelines, we were in a position to explore the scientific question of which organismal traits are correlated with organelle gene counts. We gathered organelle protein-coding gene counts from the automated analysis in (Giannakis et al. 2022), ORF counts from NCBI RefSeq (O’Leary et al. 2016), and organismal properties from the data sources above, comprising 67 ecological and organismal features (Methods; Supplementary Tables 1–2). We used estimated taxonomic trees from NCBI’s Common Taxonomy Tree Tool (Federhen 2012). For each property, we considered each possible value in turn as the “positive” case (an illustration, for the parasitism trait, is shown in Fig. 4). The dataset showed strong phylogenetic signal, with Pagel’s λ exceeding 0.999 for both organelle counts.

**Figure 4.**
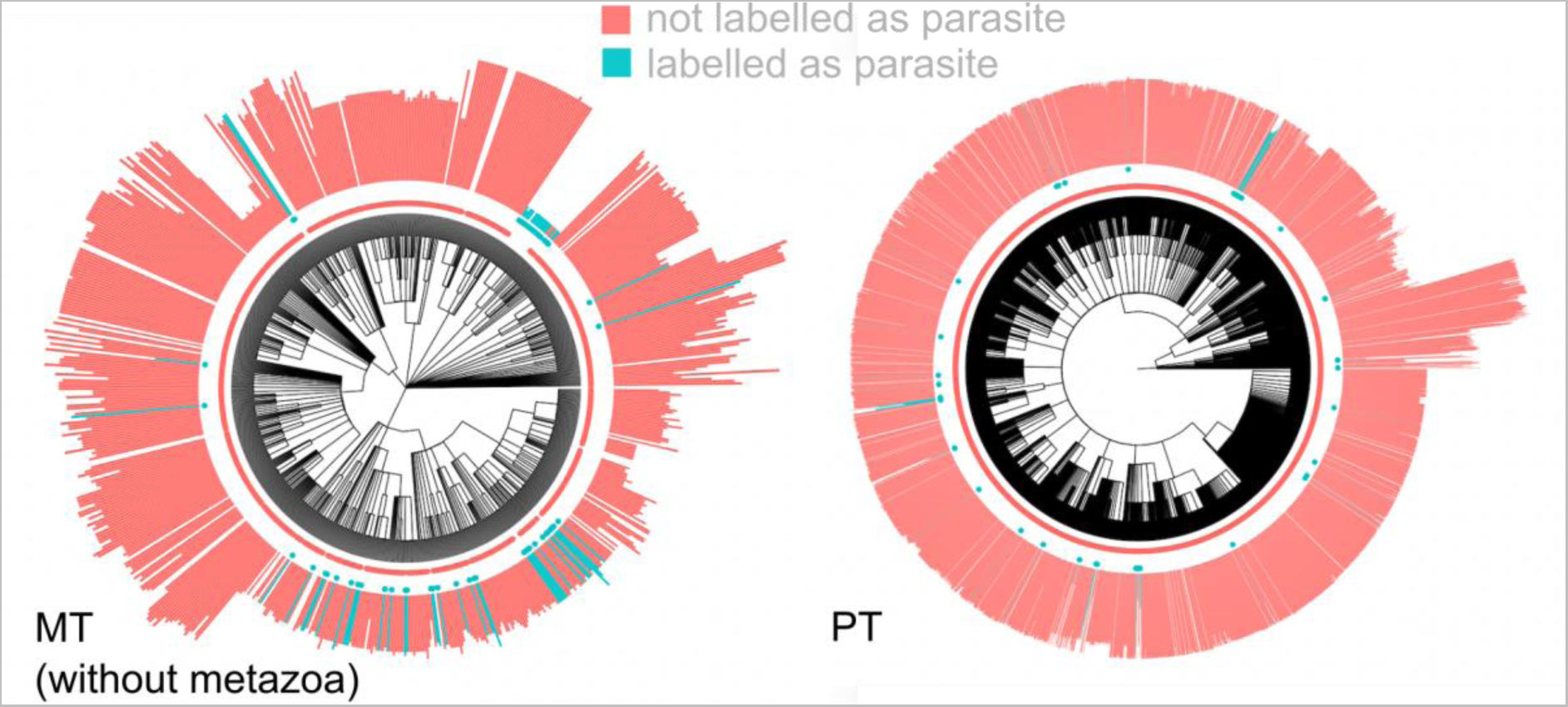
Example oDNA data. (left) mtDNA, without metazoa for clarity; (right) ptDNA. Colour (and central point markers) give the predictor value (parasitism positive, blue, or negative, red). Length of bars gives oDNA gene count.

Several issues immediately arose in the comparative analysis of these real datasets. While our benchmarking approach tested the effect of traits not being measured for all individuals, in the ecological data, not all traits are even defined for all eukaryotes. Initial empirical investigation suggested, for example, a link between the feature “palm” and mtDNA gene count. But further investigation quickly showed that this was simply reporting the fact that palms are plants and animals, which dominate the dataset, are not. In this and many other cases, the predictor can be regarded not just as negative, but undefined, for a subset of species. In these cases, we perform analysis only on the subtree rooted at the common ancestor of all positive-labelled species (thus restricting the “palm” analysis to plants). We used both PLM with a Brownian-motion derived correlation structure (reasoning that the strong agreement between PLM and PGLS results justified the choice of the much faster approach) and PGLM to explore links between the (predictor) property and the (gene count) response.

The next issue was one of (outlier) influence. In several cases, a significant correlation was reported by the pipeline but was completely dependent on a single observation. An example is given in Supplementary Fig. 5, where the “effect” arises from a shifted ratio of *n*=12 to *n*=13 genes in Metazoa, but the ratio in the positive case is highly dependent on a single *n*=12 observation. To avoid dubious cases like this, we filtered results: if either predictor value had only two associated response values, we required at least six observations of all response values to retain a result.

Finally, we found some issues in the original dataset, including, for example, metazoans being assigned plant traits. We identified and fixed these issues manually. The results for predictors of confirmed protein-coding gene counts, and CDS region counts, are shown in Fig. 5 and summarized in Table 1. We report both results that are robust to Bonferroni correction respecting all the predictors tested, and those that display p < 0.05 but are not robust to Bonferroni correction (which could be viewed as somewhat conservative, especially given the large number of predictors considered).

**Figure 5.**
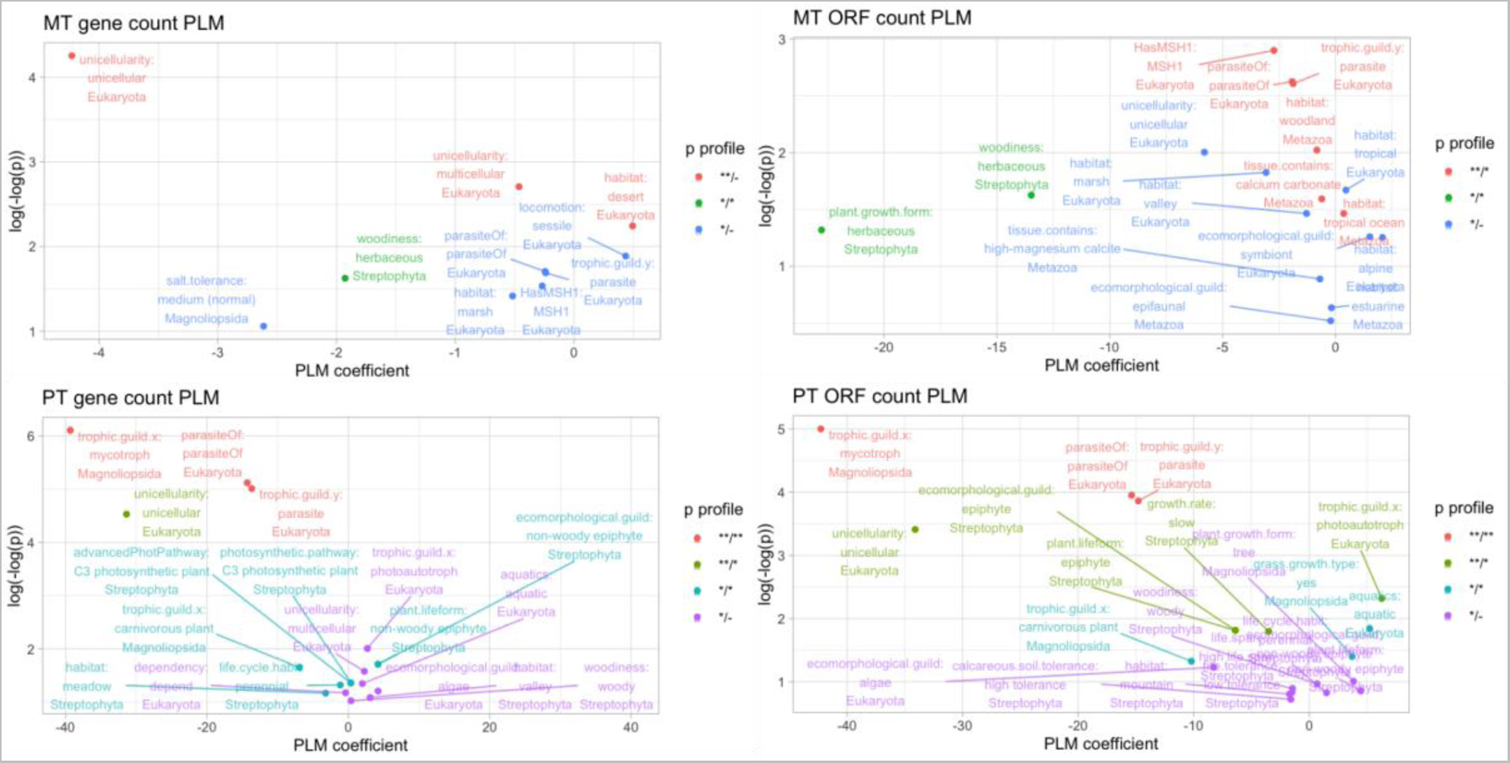
Features correlated with oDNA gene counts. PLM coefficients (x-axis) and p-values (y-axis, double-logged and inverted) for relationships between different organismal traits and organelle DNA gene counts (mtDNA and ptDNA), counted as confirmed protein-coding genes or CDS regions. This analysis is applied to the cross-eukaryote dataset as described in the text. Colours correspond to profiles of statistical significance using PLM and PGLM approaches; ** denotes p < 0.05 after Bonferroni; * p < 0.05 without correction; - p > 0.05 (for example, **/* means one approach gave a Bonferroni-robust p < 0.05 and the other gave 0.05 not robust to Bonferroni). The PLM coefficient gives the average inferred change in gene count if an organism has a given property. The majority of traits give substantially higher p-values and lower-magnitude coefficients; plots are vertically truncated to focus on the more robust results. Example of the full distributions of oDNA gene counts with different features can be seen in (Supplementary Figs. 8–9).

**Table 1.** Summary of identified predictors of oDNA gene counts. Direction gives the sign of the coefficient of a predictor on gene count response. As in Fig. 5, p-value profiles describe statistical significance in PLM and PGLM approaches: ** denotes p < 0.05 after Bonferroni; * p < 0.05 without correction; - p > 0.05. For example, parasitism is correlated with mtDNA gene count at the p < 0.05 level by one of PGLM or PLM (*/-), and with mtDNA ORF count at the Bonferroni p < 0.05 level by one approach and the p < 0.05 level by the other (**/*). † both unicellularity and multicellularity appear with gene count effects in the same direction – reflecting a probably artefactual comparison with the unlabelled subset of the data. †† non-woody epiphytes appear as a significant correlate, but with the opposite sign, in the gene count dataset, suggesting an interaction between woodiness and epiphyte traits.

| Feature | Clade | Direction | Gene count p-val profile | ORF count p-val profile |
| --- | --- | --- | --- | --- |
| **mtDNA** |  |  |  |  |
| Parasitism | Eukaryota | - | */- | **/* |
| Herbaceous growth form | Streptophyta | - | */* | */* |
| Woodland habitat | Metazoa | - | -/- | **/* |
| Tropical ocean habitat | Metazoa | + | -/- | **/* |
| Has *msh1* | Eukaryota | - | */- | **/* |
| CaCO<sub>3</sub> tissue | Metazoa | + | -/- | **/* |
| Unicellularity/multicellularity† | Eukaryota | - | **/- | */- |
| Desert habitat | Eukaryota | + | **/- | -/- |
| **ptDNA** |  |  |  |  |
| Parasitism | Eukaryota | - | **/** | **/** |
| Mycotroph | Magnoliopsida | - | **/** | **/** |
| Unicellularity | Eukaryota | - | **/* | **/* |
| Epiphyte†† | Streptophyta | - | -/- | **/* |
| Photoautotroph | Eukaryota | + | */- | **/* |
| Slow growth rate | Streptophyta | - | -/- | **/* |
| Carnivorous plant | Magnoliopsida | - | */* | */* |
| Aquatic | Eukaryota | + | */- | */* |

We also conducted the same analysis for two alternative phylogenies, in an attempt to further improve transparency. In case the oversampling of metazoan mtDNA retained an influence on results from other taxa, we also removed metazoans from the mtDNA data and performed the same analysis (Supplementary Fig. 6); most of the non-metazoan traits in Fig. 5 and Table 1 were retained, and an additional significant link with a tropical habitat (associated with higher mtDNA gene counts) was observed. Finally, we also performed the analysis with a smaller, plants-only phylogeny with complete branch length estimates (also removing these estimates to further test the influence of branch length information), in Supplementary Fig. 7. Here, the ptDNA correlates were very comparable to those from the cross-eukaryotic data, but the mtDNA correlates differed. Desert habitat and herbaceous plant form were retained, but several features relating to environmental tolerance (soil and shade, for example) also appeared. The results from neglecting branch lengths in this plant dataset were largely comparable to those retaining branch length estimates, but with less statistical power (typically less robust p-values).

Our approaches blocking, or assigning random effects to, eukaryotic clade and analysing residual variance (see Methods) broadly agreed with these features. For the non-parameteric approach blocking clade (Supplementary Figs. 8–9), the set of suggested predictor factors at the Bonferroni p < 0.05 level for mtDNA including traits related to parasitism, unicellularity, plant lifeform, and animal motility. For ptDNA the traits were parasitism, plant growth form and lifeform, carnivory and mycotrophy, unicellularity, and leaf morphology. The mixed-model approaches assigning random effects to clade, and the approach where gene counts were normalized by subtracting the clade’s mean count, produced broadly similar results to Fig. 5 (Supplementary Figs. 10–11). Here, parasitism and unicellularity appear once more for mtDNA gene counts, with herbaceous plant form, sessility, bryophytic lifeform, and forest environments also appearing. Some further features identified by the mixed-model approach alone include C_3_ photosynthesis and broad leaf morphology (both associated with lower mtDNA gene counts) among Streptophyta. For ptDNA the mixed modelling approach once more identified parasitism, mycotrophy, carnivory, and unicellularity, with some further features including a meadow habitat and perennial life cycle (both associated with lower ptDNA gene counts). It must be repeated that these approaches only account for relatedness at the highest level of the eukaryotic tree, but do attempt to directly control for the sometimes dramatic systematic differences in oDNA gene counts across different clades.

## Discussion

For both mitochondria and chloroplasts, we anticipated one known effect: parasitic species often retain fewer organelle DNA genes. However, particularly in the case of mtDNA, it was not clear how strong the associated signal would be: many parasitic animals exist, for example, that have the same mtDNA gene profile as free-living animals. Many deep-branching taxa, including Apicomplexans, are parasitic and have highly reduced (or no) mtDNA – but such examples may reflect similarity by descent and in fact provide little support for a general relationship across life (Fig. 4). However, parasitism was readily detected by the pipeline (Fig. 5, Table 1), and in the ptDNA case, several other features related to a reduced dependence on photosynthesis (mycotrophy, carnivory) clearly emerged as predictors of reduced ptDNA counts.

It has been hypothesized that organelle gene retention may in part depend on features of an organism’s lifestyle and environment (Johnston 2019; García-Pascual, Nordbotten, and Johnston 2022); an instance of the principle of colocalization for redox regulation (Allen 2015; Allen and Martin 2016). The picture here is that if an environment imposes strong, varying demands on an organism’s metabolism and bioenergetic budget, the retention of oDNA genes may help organelles rapidly and individually respond to these challenges. By contrast, organisms in more stable, less challenging environments can allow organelle genes to transfer to the nucleus, as rapid response is less important and the nucleus provides a genetically preferable environment.

Several of the ecological traits suggested by this analysis support this picture. PtDNA counts are lower in organisms that are less exposed to photosynthetic demands: parasites and those species with other nutrient and energy sources (mycotrophs, carnivorous plants). Aquatic phototrophs and algae (a weaker signal for the latter set), both of which exist in tumultuous shallow or tidal environments, retain more ptDNA genes (García-Pascual, Nordbotten, and Johnston 2022; Giannakis et al. 2022; Keeling 2010; Mohanta et al. 2020). Parasites retain fewer mtDNA genes (even after accounting for phylogenetic links) (Hjort et al. 2010), and sessile organisms – which cannot move to evade environmental changes and challenges (Johnston 2019) – retain more. In mtDNA, there are some habitat-specific signals that also correspond with this picture: desert and tropical ocean habitats (with strong diurnal variability in temperature) are correlated with higher mtDNA gene counts, while woodland habitats (in a sense more buffered and stable) are correlated with lower ones. Weaker signals, only detected in a subset of analyses, also appear: alpine and tropical environments correlate with higher gene counts.

Other identified links fall less clearly into this broad picture, and constitute potentially interesting lines for further investigation. Herbaceous plants (vascular plants with no woody stems) and epiphytes (plants that live on other plants but which do not derive nutrients from them) have lower mtDNA and ptDNA counts respectively; metazoans with CaCO_3_ tissues seem to have a skew towards higher gene counts (inasmuch as diversity exists in metazoan mtDNA). We can very cautiously speculate a little on possible connections here. Herbaceous plants typically have shorter lifespans than woody ones, meaning that they may be exposed to comparatively little environmental change over their individual lives – potentially shifting the balance away from oDNA gene retention. Epiphytes, often protected by forest canopy and colonizing environmental niches that few other species compete for, may broadly experience more stable environments than free-standing species, again shifting the balance away from oDNA gene retention. The other signals identified by individual methods, including other habitats and particularly perennial life cycles, may be interesting targets for further investigation of other mechanisms, perhaps analogous to elevated organelle gene transfer in plants with selfing or clonal reproductive behaviour (Brandvain, Barker, and Wade 2007; Brandvain and Wade 2009).

It is worth mentioning one absent result. Previous theory (García-Pascual, Nordbotten, and Johnston 2022) discussed that organisms in intertidal environments – exposed to strong, regular oscillations of temperature, water, salt, and more – may experience strong pressure to retain oDNA genes (and maintain their integrity (Edwards et al. 2021)). This is qualitatively observed in, for example, plastid gene counts in seaweeds (Mohanta et al. 2020; Keeling 2010; García-Pascual, Nordbotten, and Johnston 2022). The analysis here did not highlight intertidal habitat as a strong predictor of oDNA gene count after accounting for phylogenetic correlations, although other habitat conditions (including desert and tropical ocean) subject to strong variability were detected. Of course, given the sparse sampling in our source data, an absence of a positive result cannot directly be interpreted as a negative one. Further, detailed investigation with a specific experimental design focused on particular environments – rather than the broad approach deliberately adopted here – will provide future refinement of this picture.

These oDNA links must certainly be interpreted with some caution, for both scientific and technical reasons. Beginning with the scientific, (Janouškovec et al. 2017) have demonstrated that oDNA gene patterns emerged rather earlier in many taxa, with ongoing evolution of gene content slowing (to the point of stalling in many animals and fungi) over time. Hypotheses linking oDNA profiles to environmental traits should therefore consider the environments faced by ancestral organisms while this process was ongoing, as well as those faced by present-day organisms which survive with these established gene profiles.

Of course, despite our efforts to preserve specificity and sensitivity, there are several more technical reasons why some of the effects we see may be statistical artefacts. If observations are not missing completely at random but are sampled in some systematic way connected to the relationship between predictor and response, the correlations we detect will be affected. While the comparative approaches we use attempt to account for the relatedness of samples, the high weighting given to metazoa in the mtDNA dataset may lead to some skewing of results if this accounting is imperfect. As the rates of oDNA evolution differ in different clades, single-valued descriptions of correlations between species may not capture all the nuances of behaviour across the eukaryotic tree (though our synthetic control studies also have this differences between rates). Further, correlation is not causation, and even if correlations do exist between these variables across eukaryotes, such correlations can only ever be indicative of interesting research directions, not definitive proof of mechanistic relationships. Further work exploring the new links we have observed here in more biological detail will be required for a full understanding of their evolutionary relevance.

We hope, however, that this work has both illustrated the general potential of comparative approaches to identify correlations when many traditional assumptions are not met and when data is (rather) unreliably observed, and signposted some interesting avenues for further inquiry in the specific field of organelle genome evolution.

## Acknowledgments

We are grateful to Ramon Diaz-Uriarte for valuable discussions, Luke Harmon and Liam Revell for their excellent reference material on comparative methods and PGLS, and Julien Duthiel and Emmanuel Paradis for helpful correspondence about PGLS. LR was supported by the BBSRC via the MIBTP Doctoral Training Scheme. This project has received funding from the European Research Council (ERC) under the European Union’s Horizon 2020 research and innovation programme (Grant agreement No. 805046 (EvoConBiO) to IGJ).

## Supplementary Information

**Supplementary Figure 1.**
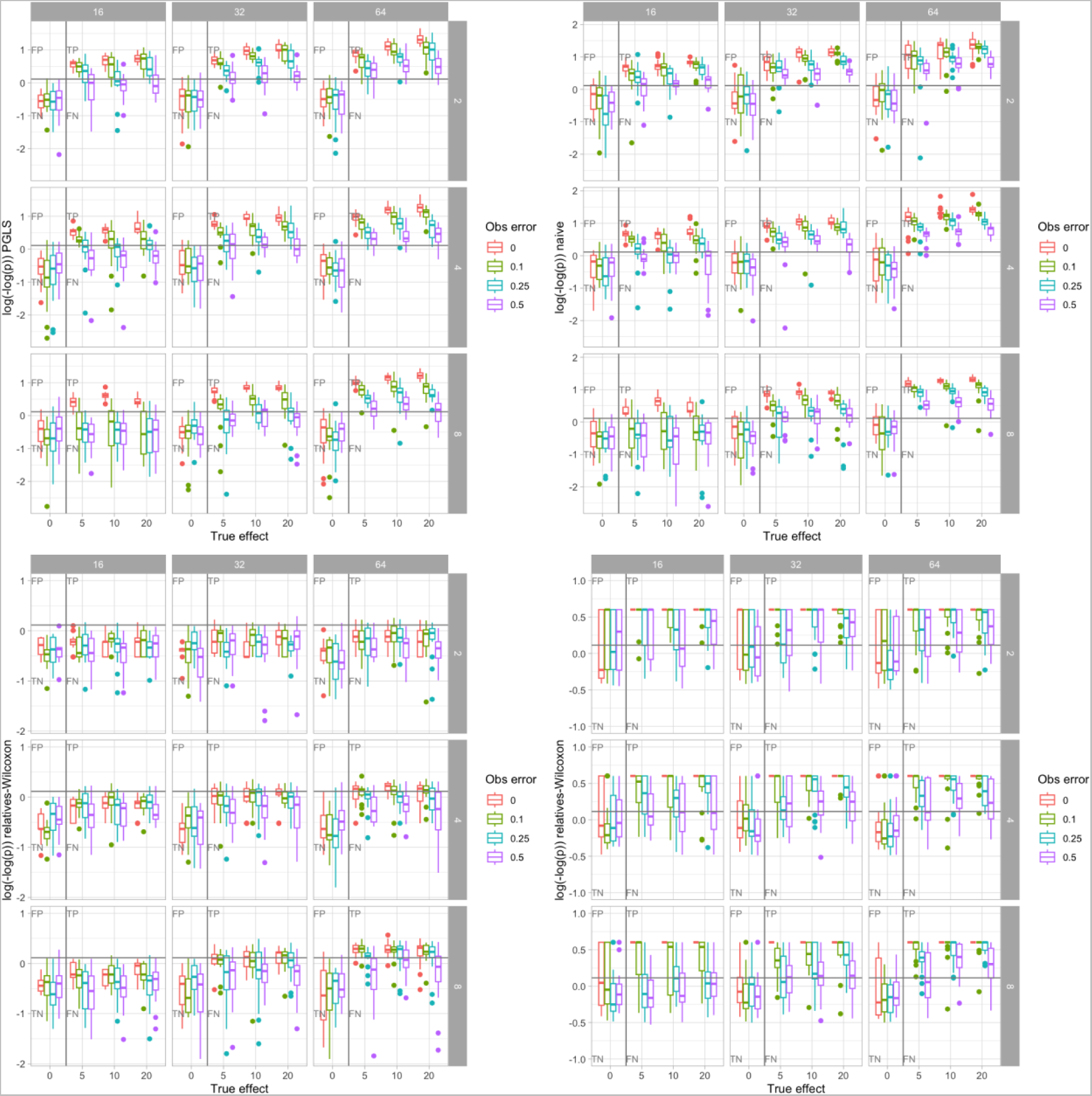
Different approaches to inferring correlations between variables in simulated taxonomically broad, sparse, unsystematized data. In each subplot, the x-axis describes *c*, the influence of the predictor on the response variable, and the y-axis gives the distribution of p-values from an analysis. On the horizontal axis, 0 is no influence (null hypothesis); increasing influence gives a stronger signal. The different colours (noise labels) correspond to different proportions of occluded observations; 0 is perfect observations of the predictor, nonzero values are the probability of observing a negative value for a positive case. The facets in each frame give the size of the tree (columns) and the average number of evolutionary events giving a positive predictor value (rows). Finally, the different frames give different inference approaches: PGLS, no phylogenetic accounting, a non-parameteric approach comparing relatives with the Mann-Whitney test, and the same principle with bootstrap resampling.

**Supplementary Figure 2.**
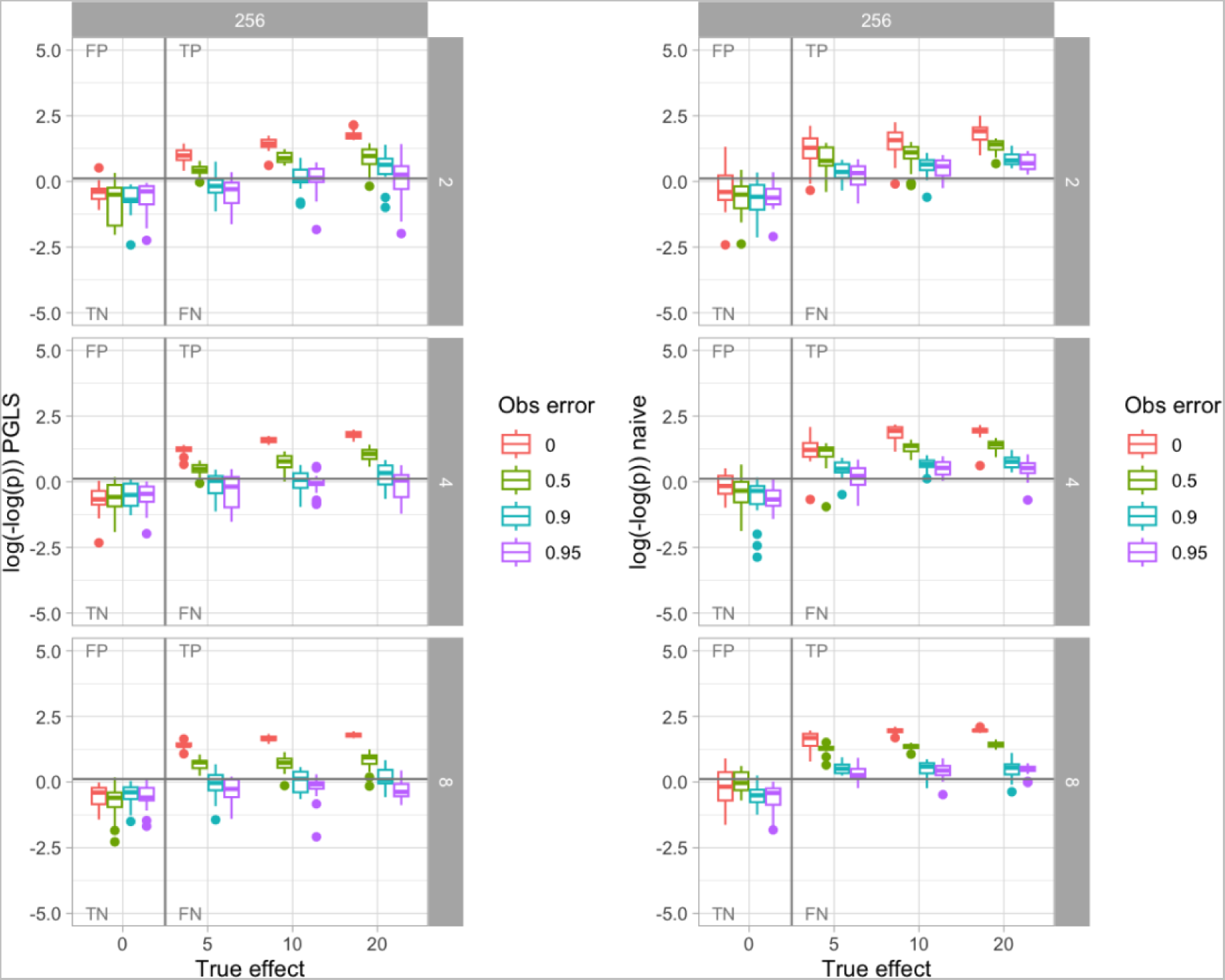
PGLS performance with changing predictor observations in simulated data. As in Supp Fig 1, the x-axis gives the strength of relationship between predictor and response, and the y-axis gives a distribution of p-values from an analysis. The average number of events generating a positive predictor value is varied across rows, and different colours give different proportions of occluded predictor observations. PGLS rarely gives false positives, and retains some statistical power even when 95% of positive predictor values are observed as negative.

**Supplementary Figure 3.**
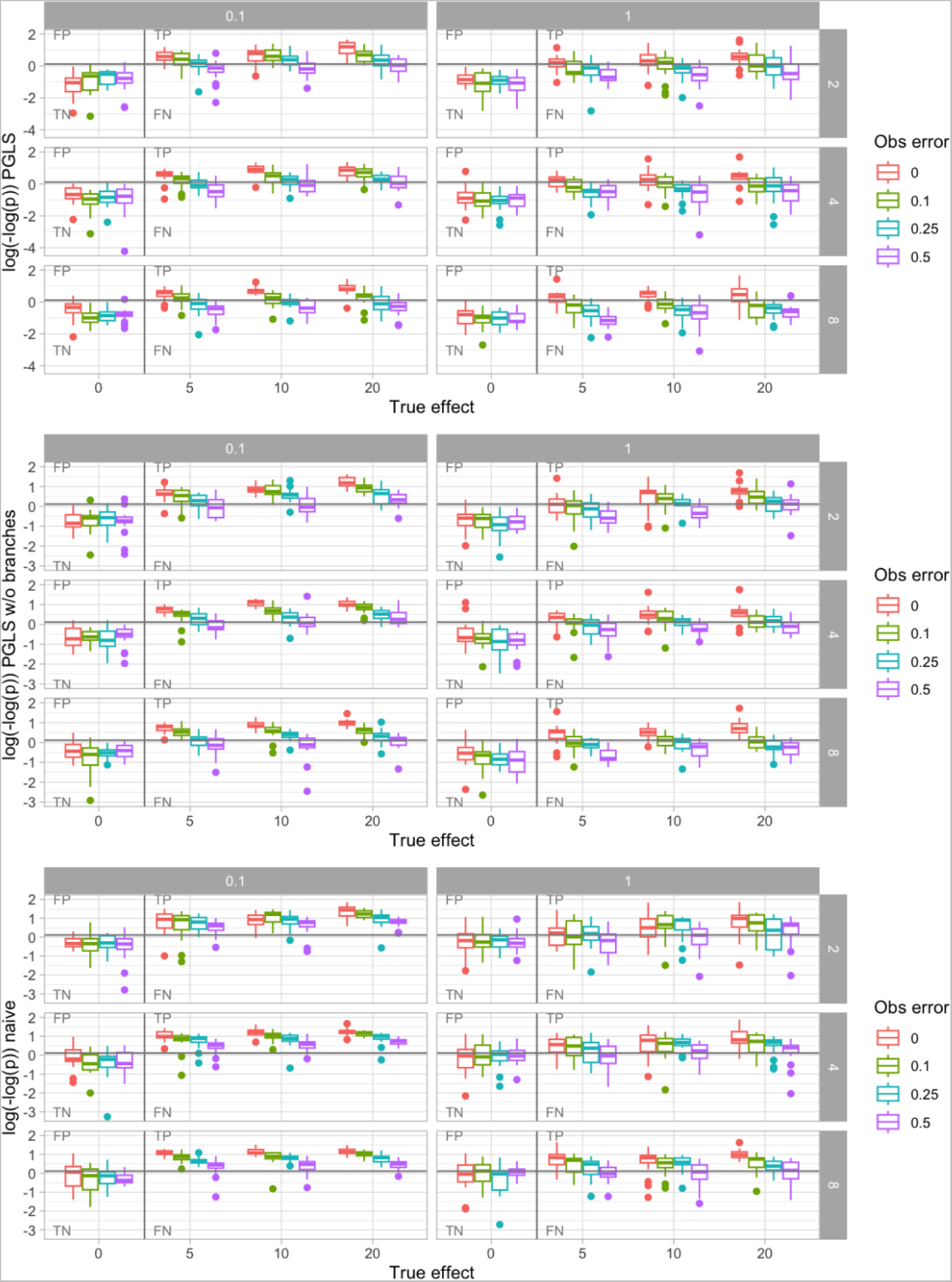
Influence of explicit branch length accounting in the inference process for simulated data. P-values and true influence of predictor on response are plotted as in Supp Fig 1 and 2. The column facets for each frame now control the death parameter in the model generating synthetic phylogenetic trees. The frames correspond to PGLS with branch length information, no phylogenetic accounting, and PGLS with uniform branch lengths (topology only).

**Supplementary Figure 4.**
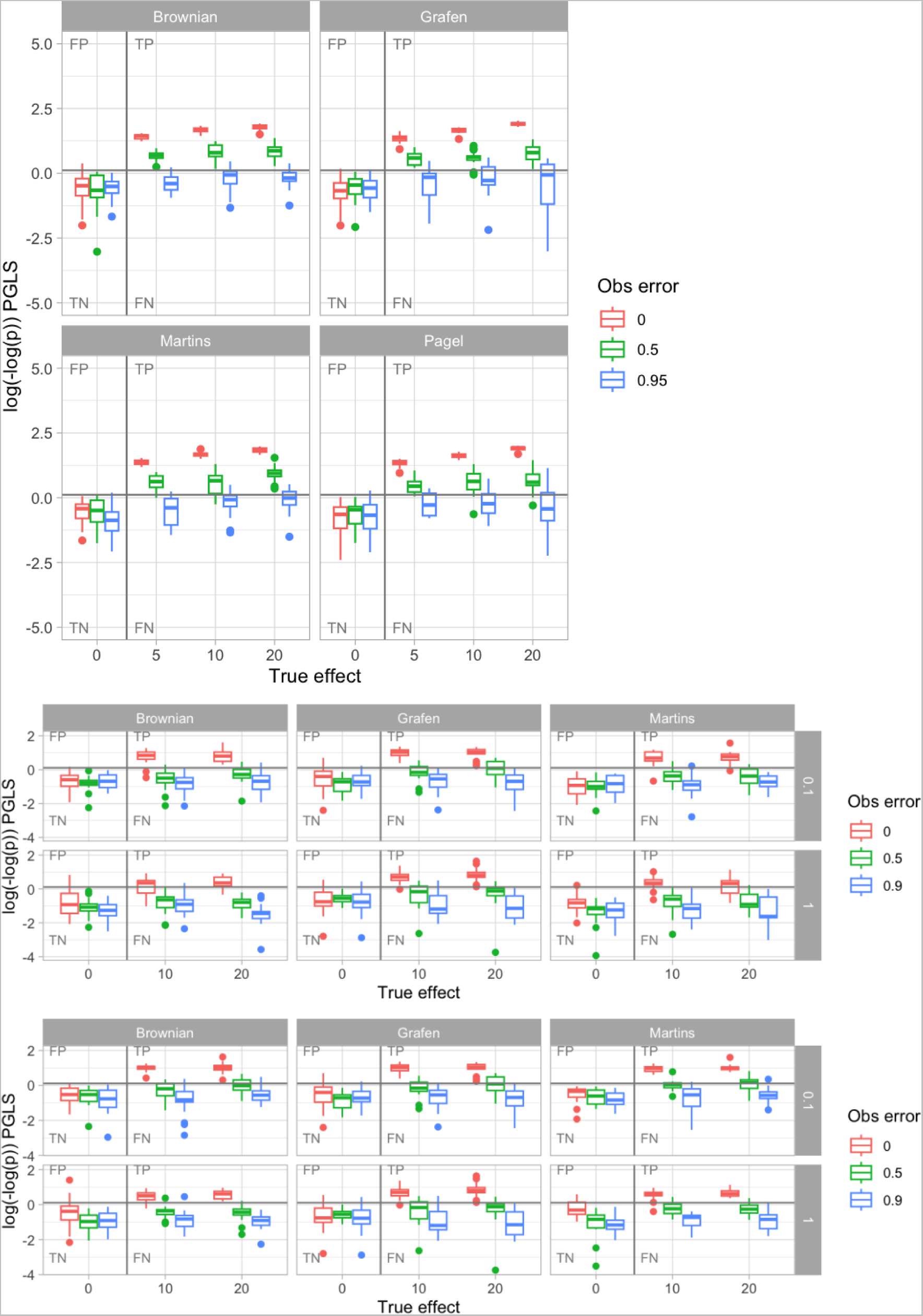
Effect of different correlation models on PGLS performance in simulated evolution. Top, symmetric tree; bottom, birth-death trees with different death parameters.

**Supplementary Figure 5.**
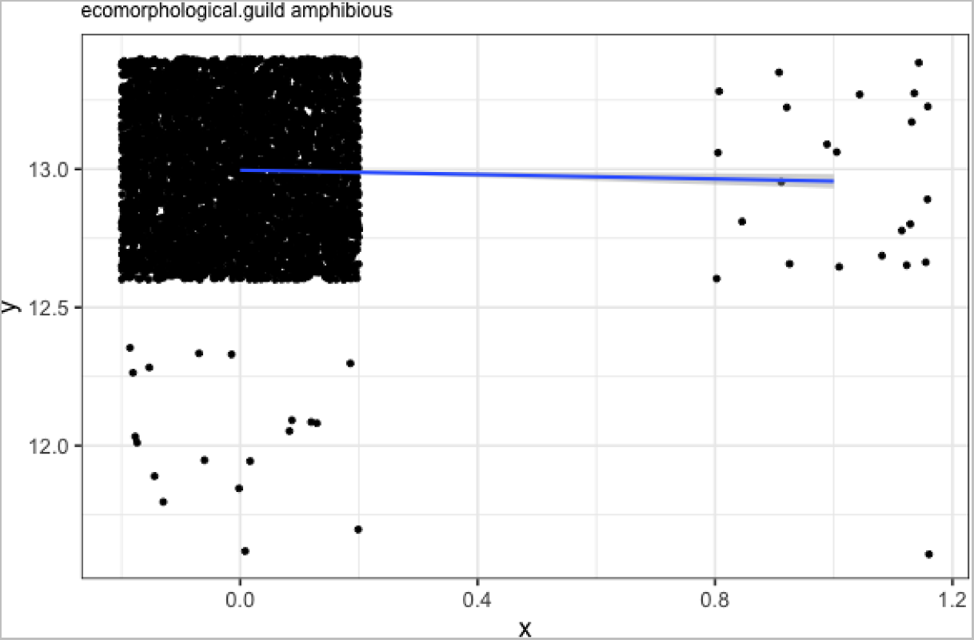
Illustration of outlier influence. Horizontal axis is predictor feature (0, not amphibious; 1, amphibious); vertical axis is gene count (12 or 13); points are jittered to show density. The vast majority of non-amphibious samples have 13 genes; the presence of the single individual with 12 genes in the much smaller amphibious sample “significantly” changes the mean. But the effect completely depends on that single observation, and therefore can likely be regarded as a sampling artefact.

**Supplementary Figure 6.**
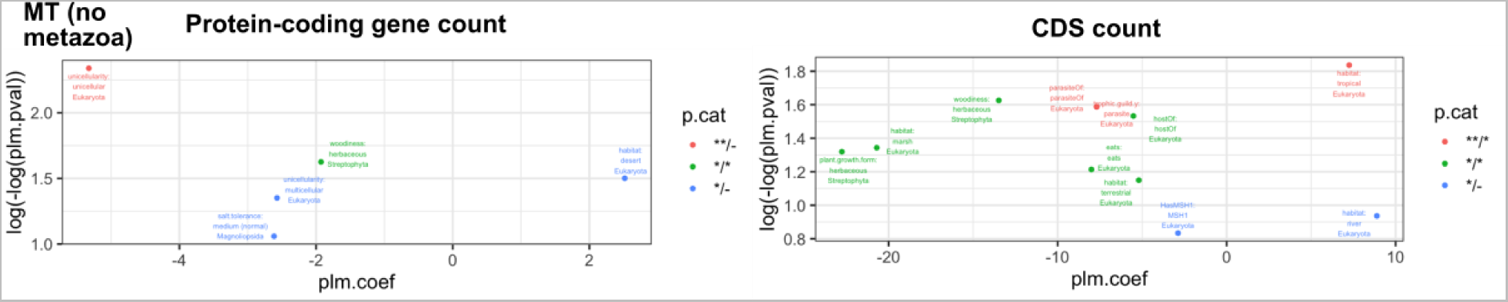
Correlations with mtDNA gene count when metazoans are removed from the phylogeny. Plots organized and labelled as in Fig. 5. PLM coefficients (x-axis) and p-values (y-axis, double-logged and inverted) for relationships between different organismal traits and mtDNA gene counts, counted as confirmed protein-coding genes or CDS regions. Colours correspond to profiles of statistical significance using PLM and PGLM approaches; ** denotes p < 0.05 after Bonferroni; * p < 0.05; - p > 0.05 (for example, **/* means one approach gave a Bonferroni-robust p < 0.05 and the other gave 0.05 not robust to Bonferroni). The PLM coefficient gives the average inferred change in gene count if an organism has a given property. The majority of traits give substantially higher p-values and lower-magnitude coefficients; plots are vertically truncated to focus on the more robust results.

**Supplementary Figure 7.**
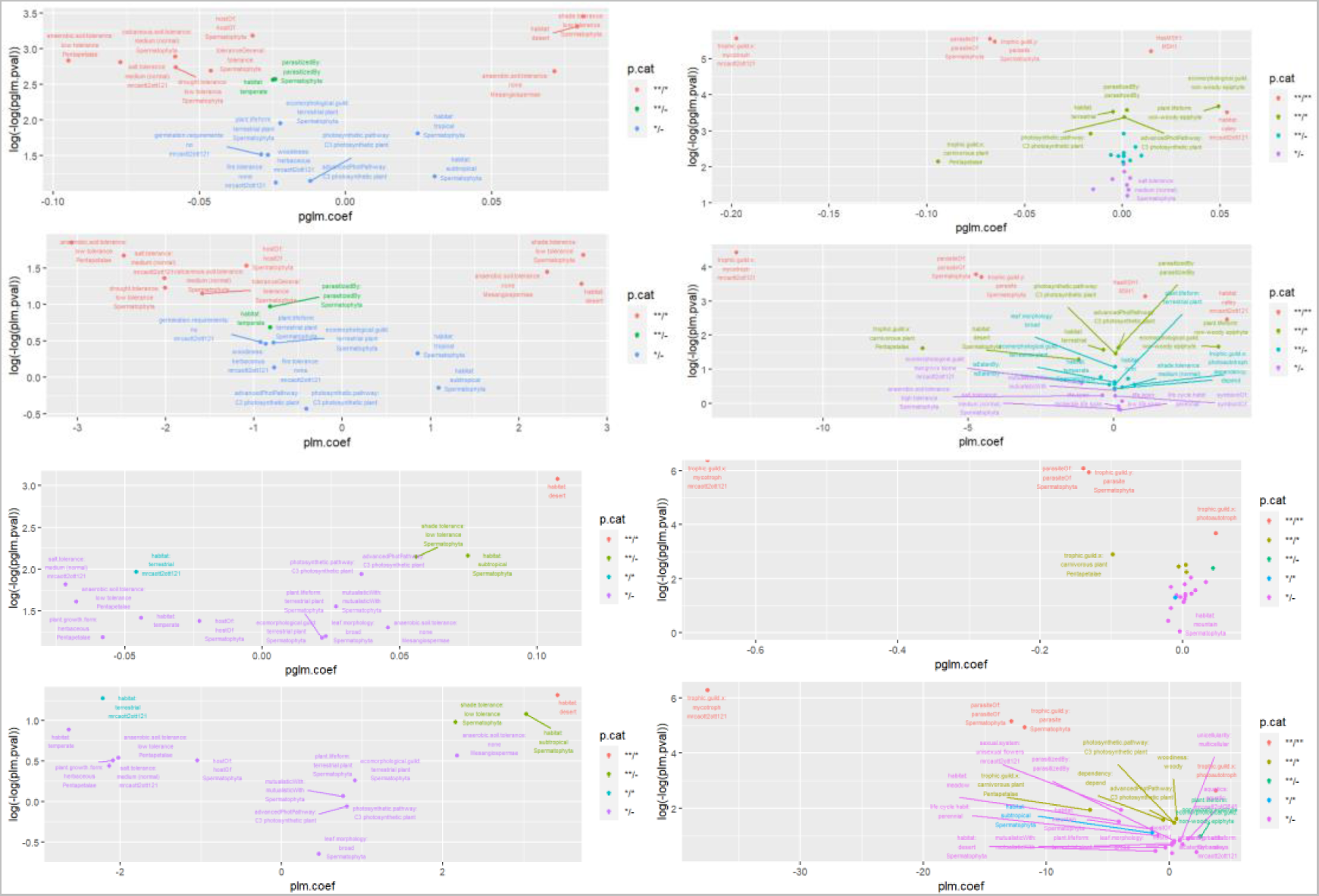
oDNA predictors in alternative plant phylogeny. Here, The plant phylogeny was adapted from the plant megaphylogeny found in (Jin and Qian 2022) comprised by phylogenies from (S. A. Smith and Brown 2018) and (Zanne et al. 2014). (left) mtDNA and (right) ptDNA correlates in plants-only phylogeny, using (top) explicit branch length estimates and (bottom) uniform branch lengths. Similar features appear as for the eukaryotic dataset for ptDNA. mtDNA connections differ (although desert habitat and herbaceouness are retained), with several features connected with soil, shade, salt, and drought tolerance appearing in these data.

**Supplementary Figure 8.**
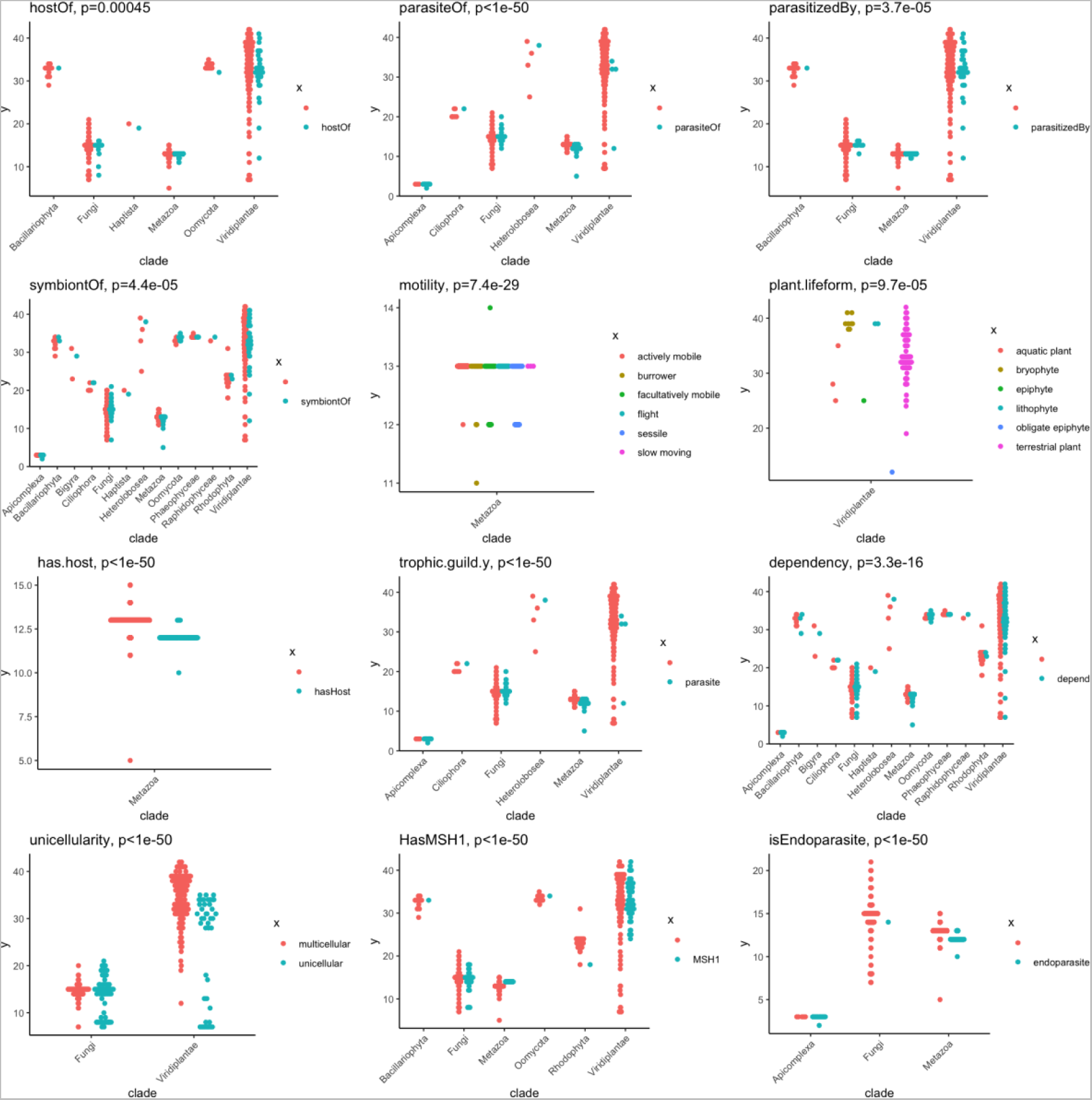
mtDNA predictors using nonparametric, non-phylogenetic approach blocking eukaryotic clade. Horizontal axis organizes samples by eukaryotic clade; colour gives different levels of predictor factor; vertical axis gives gene count. Kruskal-Wallis or Scheirer-Ray-Hare tests were used to compare gene counts with different levels of the predictor factor, treating clade as a blocking factor where appropriate.

**Supplementary Figure 9.**
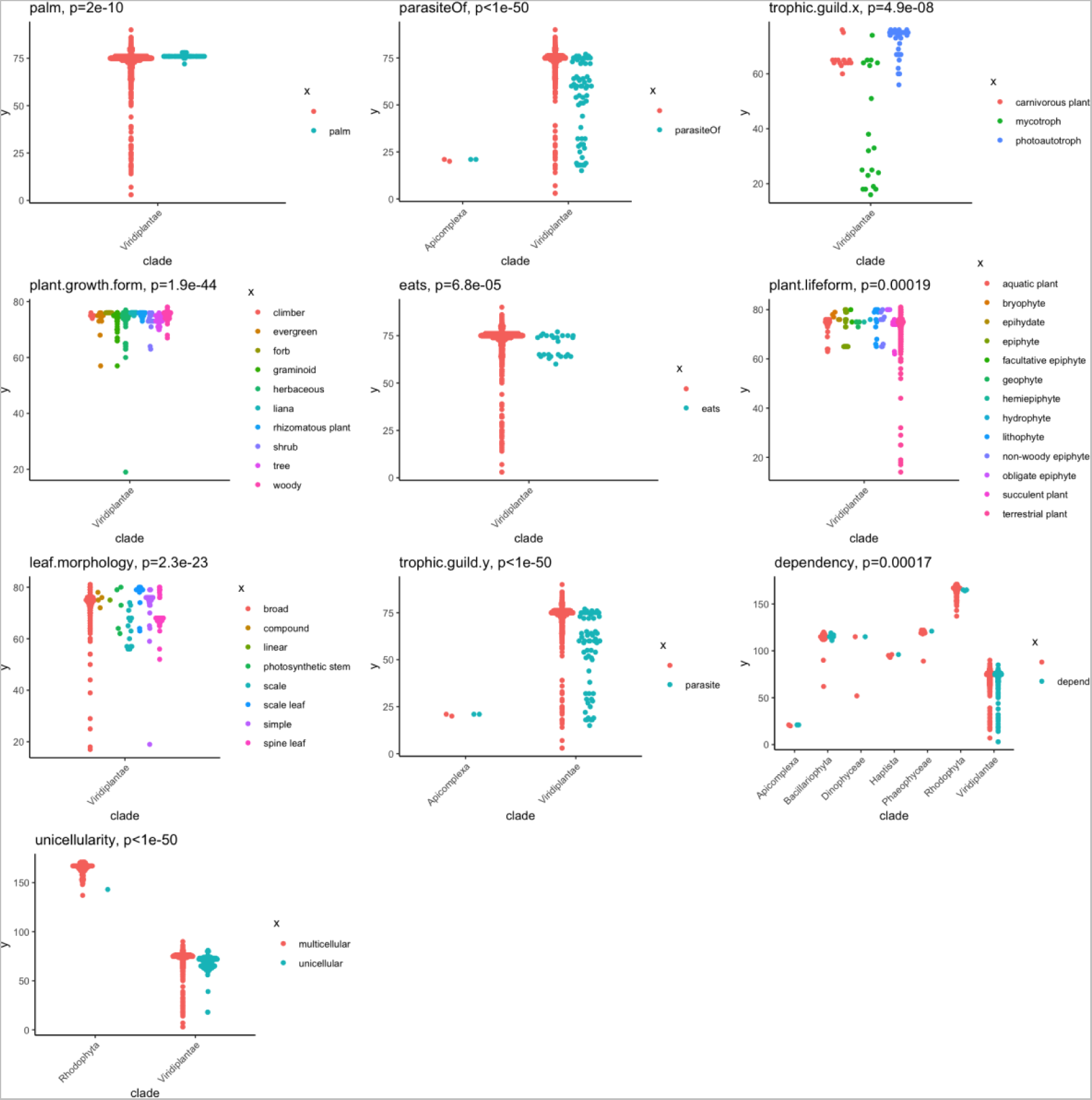
ptDNA predictors using nonparametric, non-phylogenetic approach blocking eukaryotic clade. Horizontal axis organizes samples by eukaryotic clade; colour gives different levels of predictor factor; vertical axis gives gene count. Kruskal-Wallis or Scheirer-Ray-Hare tests were used to compare gene counts with different levels of the predictor factor, treating clade as a blocking factor where appropriate.

**Supplementary Figure 10.**
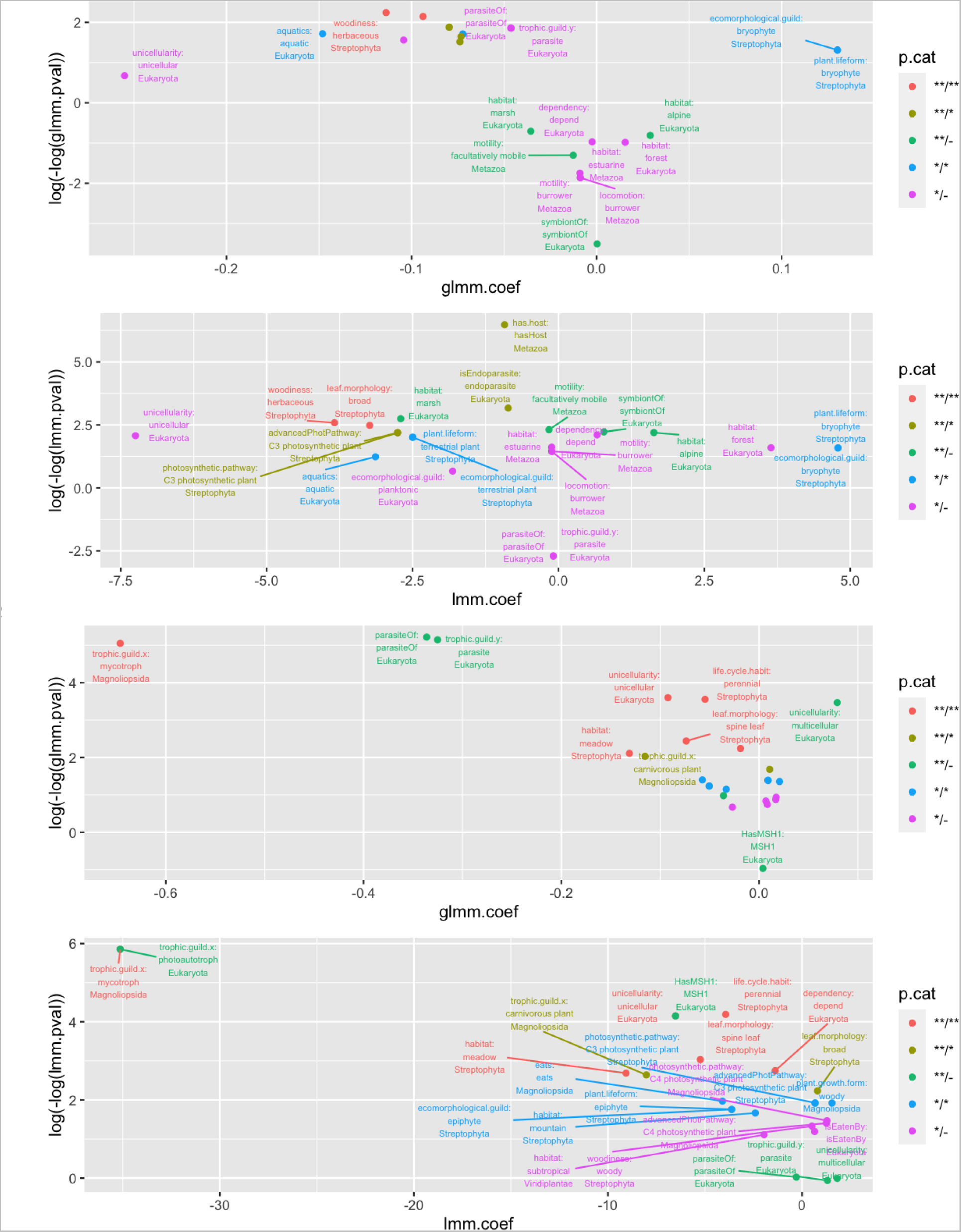
Mixed-model approaches assigning random effects to clade and analysing the remaining relationships. (top) mtDNA and (bottom) ptDNA plots as in Supplementary Fig. 7 of coefficient and p-value for linear mixed model (LMM) and Poisson generalized linear mixed model (GLMM) with random effects associated with clade, applied to either intercepts or intercepts and slopes (selected by AIC).

**Supplementary Figure 11.**
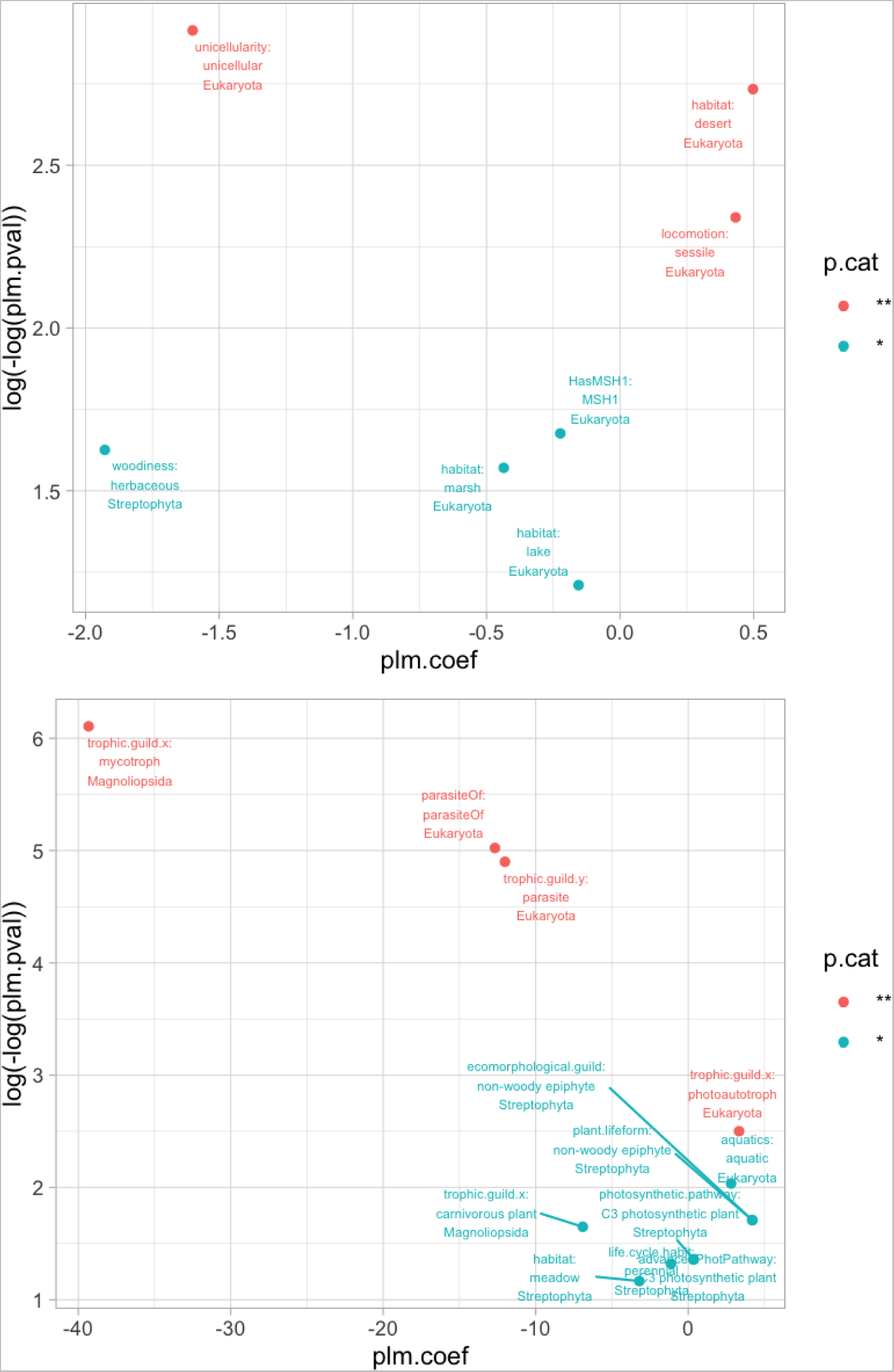
Phylogenetic linear model results after normalizing clade oDNA gene counts. The mean oDNA gene count in each clade is subtracted from all the members of that clade, then PLM is run as before, with results for mtDNA (top) and ptDNA (bottom) reported as in Fig. 5.

**Supplementary Table 1.**
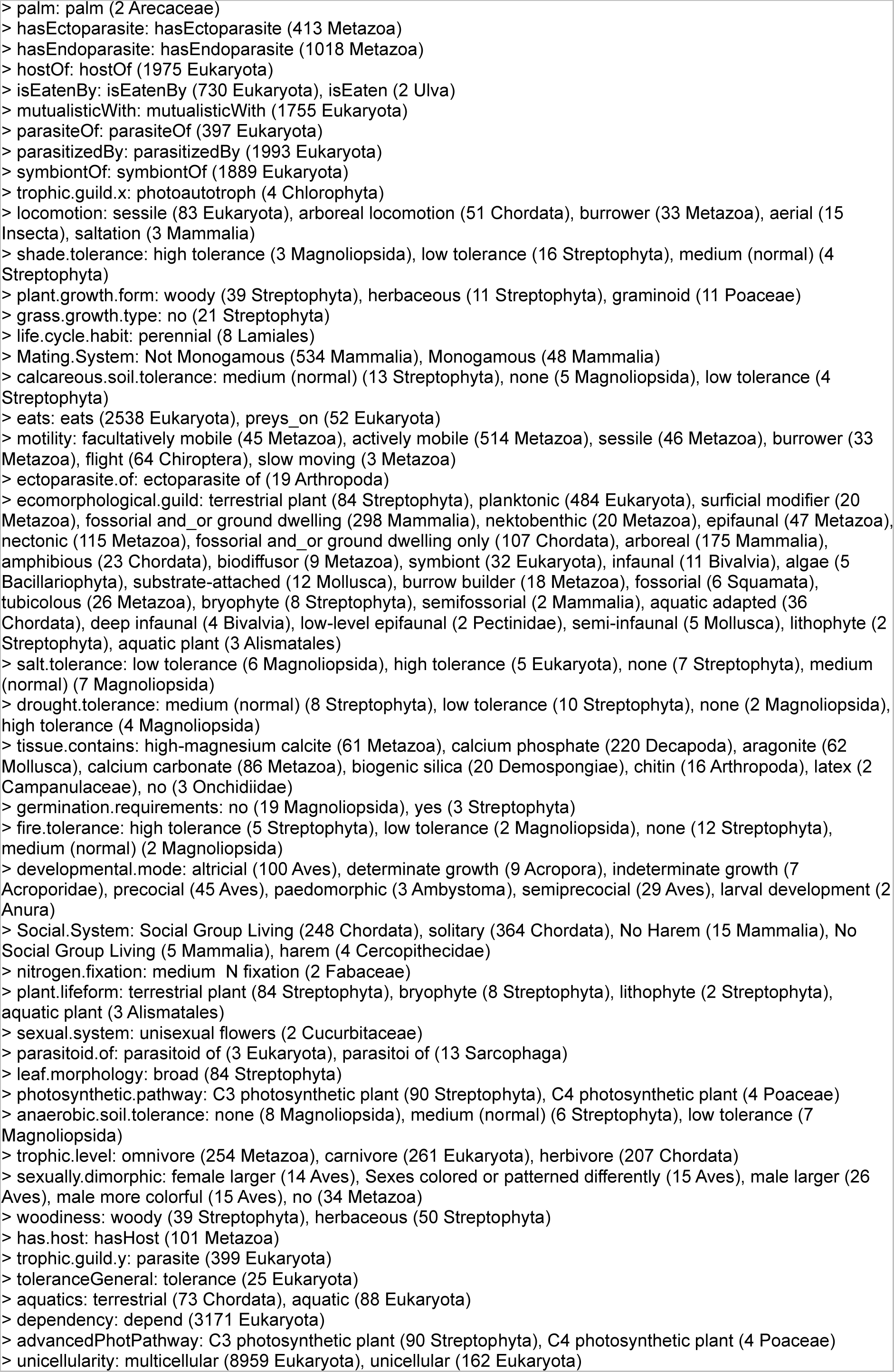
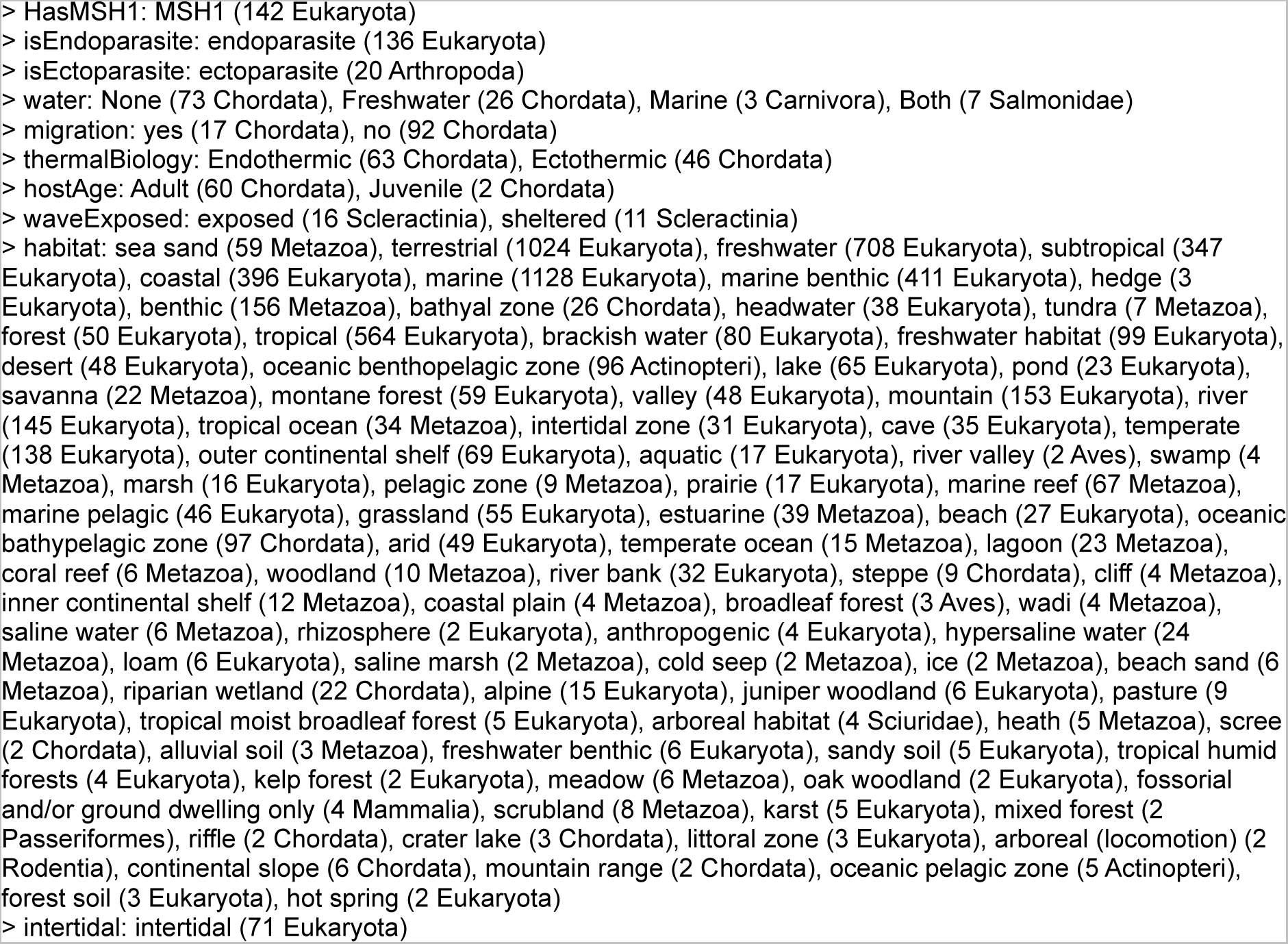
Mitochondrial ecological predictors in our compiled database. Each row gives the name of the ecological factor and each non-empty level it takes in our database of species with mitochondrial genome sequences. Each factor level is accompanied by the number of positive matches in the database and the lowest taxonomic level beneath which these matches occur.

**Supplementary Table 2.**
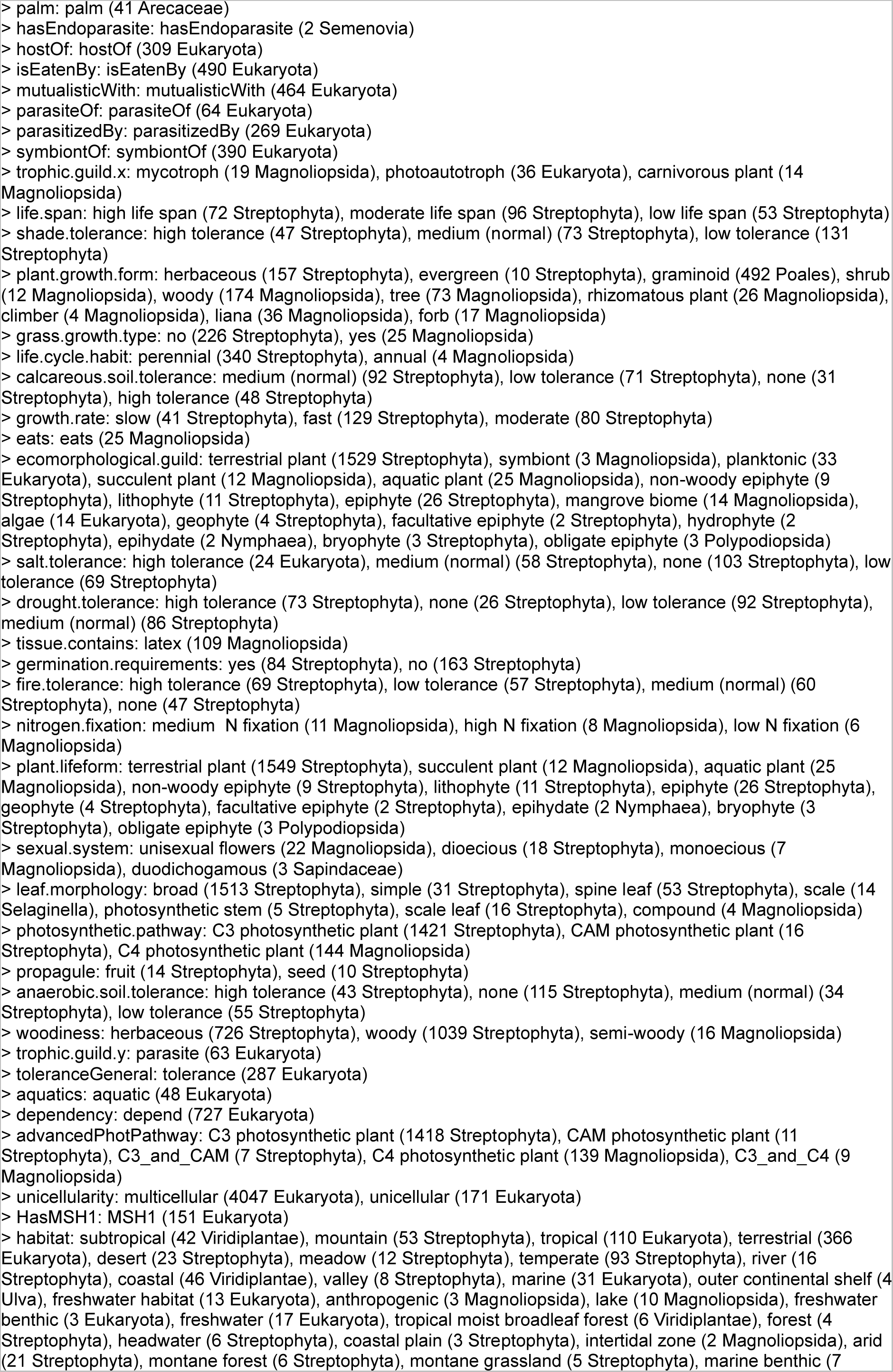
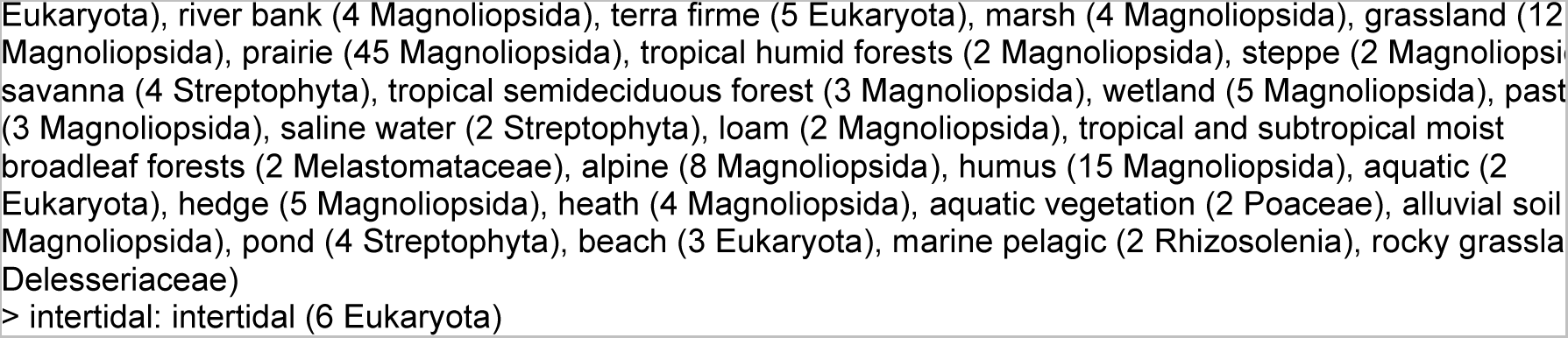
Plastid ecological predictors in our compiled database. Each row gives the name of the ecological factor and each non-empty level it takes in our database of species with plastid genome sequences. Each factor level is accompanied by the number of positive matches in the database and the lowest taxonomic level beneath which these matches occur.

## References

Allen, John F. 2015. ‘Why Chloroplasts and Mitochondria Retain Their Own Genomes and Genetic Systems: Colocation for Redox Regulation of Gene Expression’. Proceedings of the National Academy of Sciences 112 (33): 10231–38. https://doi.org/10.1073/pnas.1500012112.

Allen, John F., and William F. Martin. 2016. ‘Why Have Organelles Retained Genomes?’ Cell Systems 2 (2): 70–72. https://doi.org/10.1016/j.cels.2016.02.007.

Auguie, Baptiste, and A. Antonov. 2017. ‘GridExtra: Miscellaneous Functions for “Grid” Graphics. R Package Version 2.3’. *Computer Software]*. https://CRAN.R-Project.Org/Package=GridExtra.

Bates, Douglas, Martin Mächler, Ben Bolker, and Steve Walker. 2015. ‘Fitting Linear Mixed-Effects Models Using Lme4’. Journal of Statistical Software 67 (October): 1–48. https://doi.org/10.18637/jss.v067.i01.

Björkholm, Patrik, Ajith Harish, Erik Hagström, Andreas M. Ernst, and Siv GE Andersson. 2015. ‘Mitochondrial Genomes Are Retained by Selective Constraints on Protein Targeting’. Proceedings of the National Academy of Sciences 112 (33): 10154–61.

Brandvain, Yaniv, Michael S. Barker, and Michael J. Wade. 2007. ‘Gene Co-Inheritance and Gene Transfer’. Science (New York, N.Y.) 315 (5819): 1685. https://doi.org/10.1126/science.1134789.

Brandvain, Yaniv, and Michael J. Wade. 2009. ‘The Functional Transfer of Genes From the Mitochondria to the Nucleus: The Effects of Selection, Mutation, Population Size and Rate of Self-Fertilization’. Genetics 182 (4): 1129–39. https://doi.org/10.1534/genetics.108.100024.

Chen, Letian, and Yao-Guang Liu. 2014. ‘Male Sterility and Fertility Restoration in Crops’. Annual Review of Plant Biology 65 (1): 579–606. https://doi.org/10.1146/annurev-arplant-050213-040119.

Cohen, Jeremy M., Erin L. Sauer, Olivia Santiago, Samuel Spencer, and Jason R. Rohr. 2020. ‘Divergent Impacts of Warming Weather on Wildlife Disease Risk across Climates’. Science (New York, N.Y.) 370 (6519): eabb1702. https://doi.org/10.1126/science.abb1702.

DebRoy, S. 2006. ‘Nlme: Linear and Nonlinear Mixed Effects Models’. R Package.

Edwards, David M., Ellen C. Røyrvik, Joanna M. Chustecki, Konstantinos Giannakis, Robert C. Glastad, Arunas L. Radzvilavicius, and Iain G. Johnston. 2021. ‘Avoiding Organelle Mutational Meltdown across Eukaryotes with or without a Germline Bottleneck’. PLOS Biology 19 (4): e3001153. https://doi.org/10.1371/journal.pbio.3001153.

Eklund, Aron. 2016. ‘Beeswarm: The Bee Swarm Plot, an Alternative to Stripchart’. R Package Version 0.2 3 (4).

Federhen, Scott. 2012. ‘The NCBI Taxonomy Database’. Nucleic Acids Research 40 (D1): D136–43. https://doi.org/10.1093/nar/gkr1178.

Felsenstein, Joseph. 1985. ‘Phylogenies and the Comparative Method’. The American Naturalist 125 (1): 1–15.

Freckleton, R. P., P. H. Harvey, and M. Pagel. 2002. ‘Phylogenetic Analysis and Comparative Data: A Test and Review of Evidence.’ The American Naturalist 160 (6): 712–26. https://doi.org/10.1086/343873.

García-Pascual, Belén, Jan M. Nordbotten, and Iain G. Johnston. 2022. ‘Cellular and Environmental Dynamics Influence Species-Specific Extents of Organelle Gene Retention’. bioRxiv. https://doi.org/10.1101/2022.10.17.512581.

Giannakis, Konstantinos, Samuel J. Arrowsmith, Luke Richards, Sara Gasparini, Joanna M. Chustecki, Ellen C. Røyrvik, and Iain G. Johnston. 2022. ‘Evolutionary Inference across Eukaryotes Identifies Universal Features Shaping Organelle Gene Retention’. Cell Systems 13 (11): 874–884.e5. https://doi.org/10.1016/j.cels.2022.08.007.

Grafen, Alan. 1989. ‘The Phylogenetic Regression’. *Philosophical Transactions of the Royal Society of London. B*, Biological Sciences 326 (1233): 119–57.

Havey, Michael J. 2004. ‘The Use of Cytoplasmic Male Sterility for Hybrid Seed Production’. In Molecular Biology and Biotechnology of Plant Organelles: Chloroplasts and Mitochondria, edited by Henry Daniell and Christine Chase, 623–34. Dordrecht: Springer Netherlands. https://doi.org/10.1007/978-1-4020-3166-3_23.

Heijne, Gunnar von. 1986. ‘Why Mitochondria Need a Genome’. FEBS Letters 198 (1): 1–4.

Hjort, Karin, Alina V. Goldberg, Anastasios D. Tsaousis, Robert P. Hirt, and T. Martin Embley. 2010. ‘Diversity and Reductive Evolution of Mitochondria among Microbial Eukaryotes’. Philosophical Transactions of the Royal Society B: Biological Sciences 365 (1541): 713–27. https://doi.org/10.1098/rstb.2009.0224.

Ives, Anthony R., and Theodore Garland Jr. 2010. ‘Phylogenetic Logistic Regression for Binary Dependent Variables’. Systematic Biology 59 (1): 9–26. https://doi.org/10.1093/sysbio/syp074.

Janouškovec, Jan, Denis V. Tikhonenkov, Fabien Burki, Alexis T. Howe, Forest L. Rohwer, Alexander P. Mylnikov, and Patrick J. Keeling. 2017. ‘A New Lineage of Eukaryotes Illuminates Early Mitochondrial Genome Reduction’. Current Biology 27 (23): 3717–3724.e5. https://doi.org/10.1016/j.cub.2017.10.051.

Jin, Yi, and Hong Qian. 2022. ‘V.PhyloMaker2: An Updated and Enlarged R Package That Can Generate Very Large Phylogenies for Vascular Plants’. Plant Diversity 44 (4): 335–39. https://doi.org/10.1016/j.pld.2022.05.005.

Jin, Yi, and Hong Qian. 2023. ‘U.PhyloMaker: An R Package That Can Generate Large Phylogenetic Trees for Plants and Animals’. Plant Diversity 45 (3): 347–52. https://doi.org/10.1016/j.pld.2022.12.007.

Johnston, Iain G. 2019. ‘Tension and Resolution: Dynamic, Evolving Populations of Organelle Genomes within Plant Cells’. Molecular Plant 12 (6): 764–83. https://doi.org/10.1016/j.molp.2018.11.002.

Johnston, Iain G., and Joerg P. Burgstaller. 2019. ‘Evolving MtDNA Populations within Cells’. Biochemical Society Transactions 47 (5): 1367–82.

Johnston, Iain G., and Ben P. Williams. 2016. ‘Evolutionary Inference across Eukaryotes Identifies Specific Pressures Favoring Mitochondrial Gene Retention’. Cell Systems 2 (2): 101–11. https://doi.org/10.1016/j.cels.2016.01.013.

Kattge, Jens, Gerhard Bönisch, Sandra Díaz, Sandra Lavorel, Iain Colin Prentice, Paul Leadley, Susanne Tautenhahn, Gijsbert DA Werner, Tuomas Aakala, and Mehdi Abedi. 2020. ‘TRY Plant Trait Database–Enhanced Coverage and Open Access’. Global Change Biology 26 (1): 119–88.

Keeling, Patrick J. 2010. ‘The Endosymbiotic Origin, Diversification and Fate of Plastids’. Philosophical Transactions of the Royal Society B: Biological Sciences 365 (1541): 729–48. https://doi.org/10.1098/rstb.2009.0103.

Kelly, Steven. 2021. ‘The Economics of Organellar Gene Loss and Endosymbiotic Gene Transfer’. Genome Biology 22 (1): 345. https://doi.org/10.1186/s13059-021-02567-w.

Kissling, W. Daniel, Henrik Balslev, William J. Baker, John Dransfield, Bastian Göldel, Jun Ying Lim, Renske E. Onstein, and Jens-Christian Svenning. 2019. ‘PalmTraits 1.0, a Species-Level Functional Trait Database of Palms Worldwide’. Scientific Data 6 (1): 178. https://doi.org/10.1038/s41597-019-0189-0.

Mackenzie, Sally A. 2010. ‘The Influence of Mitochondrial Genetics on Crop Breeding Strategies’. Plant Breeding Reviews: Wiley, 115–38.

Maddison, Wayne P. 2000. ‘Testing Character Correlation Using Pairwise Comparisons on a Phylogeny’. Journal of Theoretical Biology 202 (3): 195–204. https://doi.org/10.1006/jtbi.1999.1050.

Maddison, Wayne P., and Richard G. FitzJohn. 2015. ‘The Unsolved Challenge to Phylogenetic Correlation Tests for Categorical Characters’. Systematic Biology 64 (1): 127–36. https://doi.org/10.1093/sysbio/syu070.

Mangiafico, Salvatore. 2020. ‘Rcompanion: Functions to Support Extension Education Program Evaluation’. R Package Version 2 (10).

Martins, Emilia P., and Thomas F. Hansen. 1997. ‘Phylogenies and the Comparative Method: A General Approach to Incorporating Phylogenetic Information into the Analysis of Interspecific Data’. The American Naturalist 149 (4): 646–67. https://doi.org/10.1086/286013.

Mohanta, Tapan Kumar, Awdhesh Kumar Mishra, Adil Khan, Abeer Hashem, Elsayed Fathi Abd_Allah, and Ahmed Al-Harrasi. 2020. ‘Gene Loss and Evolution of the Plastome’. Genes 11 (10): 1133. https://doi.org/10.3390/genes11101133.

O’Leary, Nuala A., Mathew W. Wright, J. Rodney Brister, Stacy Ciufo, Diana Haddad, Rich McVeigh, Bhanu Rajput, et al. 2016. ‘Reference Sequence (RefSeq) Database at NCBI: Current Status, Taxonomic Expansion, and Functional Annotation’. Nucleic Acids Research 44 (D1): D733–45. https://doi.org/10.1093/nar/gkv1189.

Pagel, Mark. 1994. ‘Detecting Correlated Evolution on Phylogenies: A General Method for the Comparative Analysis of Discrete Characters’. Proceedings of the Royal Society of London. Series B: Biological Sciences 255 (1342): 37–45.

Pagel, Mark. 1999. ‘Inferring the Historical Patterns of Biological Evolution’. Nature 401 (6756): 877–84. https://doi.org/10.1038/44766.

Paradis, Emmanuel. 2014. ‘An Introduction to the Phylogenetic Comparative Method’. Modern Phylogenetic Comparative Methods and Their Application in Evolutionary Biology: Concepts and Practice, 3–18.

Paradis, Emmanuel, and Julien Claude. 2002. ‘Analysis of Comparative Data Using Generalized Estimating Equations’. Journal of Theoretical Biology 218 (2): 175–85. https://doi.org/10.1006/jtbi.2002.3066.

Paradis, Emmanuel, and Klaus Schliep. 2019. ‘Ape 5.0: An Environment for Modern Phylogenetics and Evolutionary Analyses in R’. Bioinformatics 35 (3): 526–28. https://doi.org/10.1093/bioinformatics/bty633.

Parr, Cynthia S., Nathan Wilson, Patrick Leary, Katja S. Schulz, Kristen Lans, Lisa Walley, Jennifer A. Hammock, et al. 2014. ‘The Encyclopedia of Life v2: Providing Global Access to Knowledge About Life on Earth’. Biodiversity Data Journal, no. 2 (April): e1079. https://doi.org/10.3897/BDJ.2.e1079.

Poelen, Jorrit H., James D. Simons, and Chris J. Mungall. 2014. ‘Global Biotic Interactions: An Open Infrastructure to Share and Analyze Species-Interaction Datasets’. Ecological Informatics 24 (November): 148–59. https://doi.org/10.1016/j.ecoinf.2014.08.005.

R Core Team, A., and R. Core Team. 2022. ‘R: A Language and Environment for Statistical Computing. R Foundation for Statistical Computing, Vienna, Austria. 2012’.

Revell, Liam J. 2010. ‘Phylogenetic Signal and Linear Regression on Species Data’. Methods in Ecology and Evolution 1 (4): 319–29. https://doi.org/10.1111/j.2041-210X.2010.00044.x.

Revell, Liam J. 2012. ‘Phytools: An R Package for Phylogenetic Comparative Biology (and Other Things)’. Methods in Ecology and Evolution, no. 2: 217–23.

Revell, Liam J., and Luke J. Harmon. 2022. Phylogenetic Comparative Methods in R. Princeton University Press.

Roger, Andrew J., Sergio A. Muñoz-Gómez, and Ryoma Kamikawa. 2017. ‘The Origin and Diversification of Mitochondria’. Current Biology: CB 27 (21): R1177–92. https://doi.org/10.1016/j.cub.2017.09.015.

Rohle, F. James. 2006. ‘A Comment on Phylogenetic Correction’. Evolution 60 (7): 1509–15. https://doi.org/10.1111/j.0014-3820.2006.tb01229.x.

Scheirer, C. James, William S. Ray, and Nathan Hare. 1976. ‘The Analysis of Ranked Data Derived from Completely Randomized Factorial Designs’. Biometrics 32 (2): 429–34. https://doi.org/10.2307/2529511.

Schliep, Klaus Peter. 2011. ‘Phangorn: Phylogenetic Analysis in R’. Bioinformatics 27 (4): 592–93. https://doi.org/10.1093/bioinformatics/btq706.

Slowikowski, K. 2021. ‘Ggrepel: Automatically Position Non-Overlapping Text Labels with’ggplot2’. R Package Version 0.9. 1, 2021’.

Smith, David Roy, and Patrick J. Keeling. 2015. ‘Mitochondrial and Plastid Genome Architecture: Reoccurring Themes, but Significant Differences at the Extremes’. Proceedings of the National Academy of Sciences 112 (33): 10177–84. https://doi.org/10.1073/pnas.1422049112.

Smith, Stephen A., and Joseph W. Brown. 2018. ‘Constructing a Broadly Inclusive Seed Plant Phylogeny’. American Journal of Botany 105 (3): 302–14. https://doi.org/10.1002/ajb2.1019.

Stadler, Tanja. 2011. ‘Simulating Trees with a Fixed Number of Extant Species’. Systematic Biology 60 (5): 676–84. https://doi.org/10.1093/sysbio/syr029.

Stewart, James B., and Patrick F. Chinnery. 2015. ‘The Dynamics of Mitochondrial DNA Heteroplasmy: Implications for Human Health and Disease’. Nature Reviews Genetics 16 (9): 530–42. https://doi.org/10.1038/nrg3966.

Susko, Edward. 2003. ‘Confidence Regions and Hypothesis Tests for Topologies Using Generalized Least Squares’. Molecular Biology and Evolution 20 (6): 862–68. https://doi.org/10.1093/molbev/msg093.

Symonds, Matthew R. E., and Simon P. Blomberg. 2014. ‘A Primer on Phylogenetic Generalised Least Squares’. In Modern Phylogenetic Comparative Methods and Their Application in Evolutionary Biology: Concepts and Practice, edited by László Zsolt Garamszegi, 105–30. Berlin, Heidelberg: Springer. https://doi.org/10.1007/978-3-662-43550-2_5.

Taylor, Robert W., and Doug M. Turnbull. 2005. ‘Mitochondrial DNA Mutations in Human Disease’. Nature Reviews Genetics 6 (5): 389–402. https://doi.org/10.1038/nrg1606.

Tung Ho, Lam si, and Cécile Ané. 2014. ‘A Linear-Time Algorithm for Gaussian and Non-Gaussian Trait Evolution Models’. Systematic Biology 63 (3): 397–408. https://doi.org/10.1093/sysbio/syu005.

Wallace, Douglas C., and Dimitra Chalkia. 2013. ‘Mitochondrial DNA Genetics and the Heteroplasmy Conundrum in Evolution and Disease’. Cold Spring Harbor Perspectives in Biology 5 (11): a021220. https://doi.org/10.1101/cshperspect.a021220.

Wickham, Hadley. 2011. ‘Ggplot2’. WIREs Computational Statistics 3 (2): 180–85. https://doi.org/10.1002/wics.147.

Xu, Shuangbin, Zehan Dai, Pingfan Guo, Xiaocong Fu, Shanshan Liu, Lang Zhou, Wenli Tang, et al. 2021. ‘GgtreeExtra: Compact Visualization of Richly Annotated Phylogenetic Data’. Molecular Biology and Evolution 38 (9): 4039–42. https://doi.org/10.1093/molbev/msab166.

Yu, Guangchuang, David K. Smith, Huachen Zhu, Yi Guan, and Tommy Tsan-Yuk Lam. 2017. ‘Ggtree: An r Package for Visualization and Annotation of Phylogenetic Trees with Their Covariates and Other Associated Data’. Methods in Ecology and Evolution 8 (1): 28–36. https://doi.org/10.1111/2041-210X.12628.

Zanne, Amy E., David C. Tank, William K. Cornwell, Jonathan M. Eastman, Stephen A. Smith, Richard G. FitzJohn, Daniel J. McGlinn, et al. 2014. ‘Three Keys to the Radiation of Angiosperms into Freezing Environments’. Nature 506 (7486): 89–92. https://doi.org/10.1038/nature12872.

